# Nonlinear effects of temperature on mosquito parasite infection across a large geographic climate gradient

**DOI:** 10.1101/2025.01.07.631804

**Authors:** Johannah E. Farner, Kelsey P. Lyberger, Lisa I. Couper, Mauricio Cruz-Loya, Erin A. Mordecai

## Abstract

Temperature drives ectothermic host – parasite interactions, making them particularly sensitive to climatic variation and change. To isolate the role of temperature, lab-based studies are increasingly used to assess and forecast disease risk under current and future climate conditions. However, in the field, the effects of temperature on parasitism may be mediated by other sources of variation, including local adaptation of hosts and parasites to one another and to the environment. To address the key knowledge gaps of how temperature influences host – parasite interactions and whether thermal responses measured in controlled experiments capture infection across temperature gradients in nature, we paired an extensive field survey of parasitism—by the ciliate *Lambornella clarki* on its tree hole mosquito host, *Aedes sierrensis*—with laboratory experiments describing parasitism thermal performance curves (TPCs) for six host populations from varying climates. We also investigated the mechanisms underlying the thermal biology of the host – parasite interaction by separately measuring TPCs for infection, host immunity, and parasite growth rates. Along the west coast of North America, across an 1100 km climate gradient spanning 12°C mean rainy season temperature variation, we found that parasitism peaked at intermediately cold temperatures of 9.2 – 10°C, and was consistent both between field seasons and with the lab experiment results. In the laboratory experiments, infection thermal responses were consistently nonlinear and peaked at 8.4 – 10°C, showing no evidence of host intraspecific variation in temperature sensitivity to parasitism. Importantly, parasitism peaked at temperatures below the thermal optimum for free-living *L. clarki* due to the balance of temperature effects on parasite growth and reproduction against the strength of the host melanization immune response. The results suggest that nonlinear responses to temperature drive parasitism in nature, and that simple lab and field studies can accurately capture the thermal biology of multilayered host – parasite interactions.

## Introduction

Climate warming is fundamentally important to infectious disease risk because temperature drives many of the constituent biological processes of host – parasite interactions, including host and parasite growth and development, contact rates, and immune functions (Kutz et al., 2005; Labaude et al., 2017; Laughton et al., 2017; Franke et al., 2019; Mordecai et al., 2019). Strong relationships between temperature and parasitism are frequently observed in laboratory experiments and captured in thermal performance curves (TPCs) that quantify how infection rates change with temperature, typically from data collected across a range of constant incubation temperatures (Angilletta 2009; Gehman et al., 2018; Kirk et al. 2018; Padfield et al., 2020). Following expectations from biochemical first principles, TPCs are commonly unimodal, with a given biological rate increasing with temperature from its critical thermal minimum, peaking at a thermal optimum, and then declining up to a critical thermal maximum (Angilletta, 2009). Researchers increasingly use TPCs to assess nonlinear shifts in disease risk based on current and forecasted climate and weather conditions (Gehman et al., 2018; Ryan et al., 2019; Aleuy et al., 2023). However, it is unclear whether TPCs measured in controlled laboratory settings fully capture the thermal response of disease in natural settings, where environmental complexity may modify or outweigh parasitism - temperature relationships (Mordecai et al., 2019; Taylor et al., 2019).

Parasitism TPCs are typically measured for single laboratory populations, in the absence of other stressors, and are infrequently compared to disease burdens along temperature gradients in the field (e.g., Gehman et al., 2018; Mordecai et al., 2019; Padfield et al., 2020). Although an increasing number of studies describe thermal responses for multiple populations or genotypes within host-parasite systems (Mazé-Guilmo et al., 2016; Ahrens et al., 2019; Franke et al., 2019; Aleuy et al., 2023; Ragonese et al., 2024), research on statistical or correlative relationships between TPCs for disease transmissibility (e.g., R_0_) and natural infection patterns has thus far relied on laboratory measurements from single host and parasite populations or clones, and field data collated from disparate sources and/or collected over small geographic areas. To date, this research has focused primarily on mosquito-borne human disease and fungal pathogens of zooplankton (Mordecai et al., 2013; 2017; Shocket, Ryan et al., 2018; Shocket, Strauss et al., 2018; Tesla et al., 2018; Shah et al., 2019; Shocket et al., 2020; Kirk et al., 2024). However, TPCs for both hosts and parasites can vary geographically within species due to local adaptation and population structure (Sinclair et al., 2012; Gaitán-Espitia et al., 2014; Mazé-Guilmo et al., 2016; Jurriaans & Hoogenboom, 2019; Aleuy et al., 2023; Couper et al., 2024), and additional environmental variability in field settings may outweigh or modify the temperature dependence of parasitism (e.g., Boots, 2000; Seppälä et al., 2008; Hall et al., 2009; Penczykowski et al., 2014; Palmer-Young et al., 2019; Brown et al., 2023). Thus, there is a critical need to address the questions of how temperature affects parasitism in the field, whether lab-based TPC estimates can accurately, and mechanistically, capture these patterns, and whether local thermal adaptation modifies the temperature-dependence of parasitism.

Central to capturing the thermal biology of parasitism is understanding the host and parasite mechanisms that drive the interaction. For example, host life history and defense traits and parasite growth, attack, dispersal, and transmission traits may each respond nonlinearly and distinctly to temperature. The combined effect of these traits determines the overall impact of temperature on parasitism (Paull et al., 2012; Shocket et al., 2018; Mordecai et al., 2019; Sun et al., 2023). Moreover, host and parasite trait thermal responses may differ across populations due to local adaptation and across environmental conditions (Thomas & Blanford, 2003; Sternberg & Thomas, 2014). Given that it is not feasible to measure TPCs for every possible combination of host population, parasite population, and environmental context, it is important to establish whether TPCs measured for a single set of populations and conditions can generalize to cover entire species ranges, and which traits capture the essential thermal biology of the interaction. Here, we address these questions in a model system of a mosquito (*Aedes sierrensis*) and its facultative ciliate parasite (*Lambornella clarki*), which occur across a large geographic gradient in western North America.

In this study, we pair field surveys along the entire west coast of the continental United States—spanning the geographic range of the host species—with laboratory experiments describing the temperature dependence of infection and key traits involved in host defenses and parasite fitness, for host populations from different climates across the survey area. Both species are short-lived ectotherms that are directly affected by environmental temperatures, occur commonly across a large geographic climate gradient in discrete tree hole habitats that are naturally replicated across landscapes, and can be reared separately and together in the lab (Washburn & Anderson, 1986; Broberg & Bradshaw, 1997; Darsie & Ward, 2005; Ismail et al., 2023; Lyberger et al., 2024). A 1986 field survey found that infection occurrence increased from southern to northern California (Washburn & Anderson, 1986). Our recent work describing temperature effects on different populations of *Ae. sierrensis, L. clarki*, and their interactions indicates that infection has a consistent nonlinear relationship with temperature for both sympatric and allopatric pairs of host and parasite populations collected across climates, despite evidence of local thermal adaptation in free-living *L. clarki* populations (Ismail et al., 2023; Lyberger et al., 2024). This body of work generates the hypothesis that parasitism responds to temperature as a function of the thermal biology of multiple host and parasite traits. However, it is unknown whether the lab-measured temperature impacts on parasitism across different *Ae. sierrensis* host populations are borne out in infection occurrence across the environmentally complex species range.

## Methods

### Study system

*Ae. sierrensis* and its facultative ciliate parasite *L. clarki* occur across western North America in seasonally rainwater-filled tree holes, where mosquitoes undergo one reproductive cycle annually in which larvae feed on detritus and microbes (including free-living *L. clarki*) before pupating and emerging as adults, and free-living *L. clarki* ciliates feed on aquatic bacteria (Washburn & Anderson, 1986; Darsie & Ward, 2005). In the presence of *Ae. sierrensis* larvae, free-living *L. clarki* irreversibly transform into parasites that attack larval mosquitoes by attaching to the larval cuticle, penetrating into the body cavity, and then replicating, often to high densities that kill hosts after a few weeks (Washburn et al., 1988). Mosquitoes can resist infection through multiple defenses that each likely respond nonlinearly to temperature: melanization (a generalized immune response), molting between larval instars, developing into pupae, and consuming free-living *L. clarki* in the water column (Clark & Brandl, 1976; Washburn et al., 1988; Washburn et al., 1991; Suwanchaichinda & Paskewitz, 1998; Christensen et al., 2005; Murdock et al., 2012; Martin & Hillyer, 2024). Beyond the previously documented latitudinal gradient, *L. clarki* presence is not correlated with habitat characteristics such as tree hole size, exposure, height above ground, or tree species (Washburn & Anderson, 1986).

### Field survey

To document infection patterns over a large geographic temperature gradient and collect materials for lab experiments, we sampled from over 500 water-filled tree hole habitats containing *Ae. sierrensis* larvae spanning 1100 km north to south with a mean November – April rainy season temperature gradient of 2.6°C to 15.3°C in California, Oregon, and Washington, USA, during the rainy seasons of 2022 and 2023 (Figure 1) (PRISM Climate Group, 2021). Each year, we started in the warmest and ended in the coolest locations to track the seasonal timing of *Ae. sierrensis* egg hatching and development (Jordan & Bradshaw, 1978; Jordan, 1980a; 1980b), collecting larval samples and examining them for infection at 10-40x magnification (Appendix S1). We aimed to sample each tree hole in both years, but had to replace some locations in 2023 due to storm damage (Figure 1).

**Figure 1.**
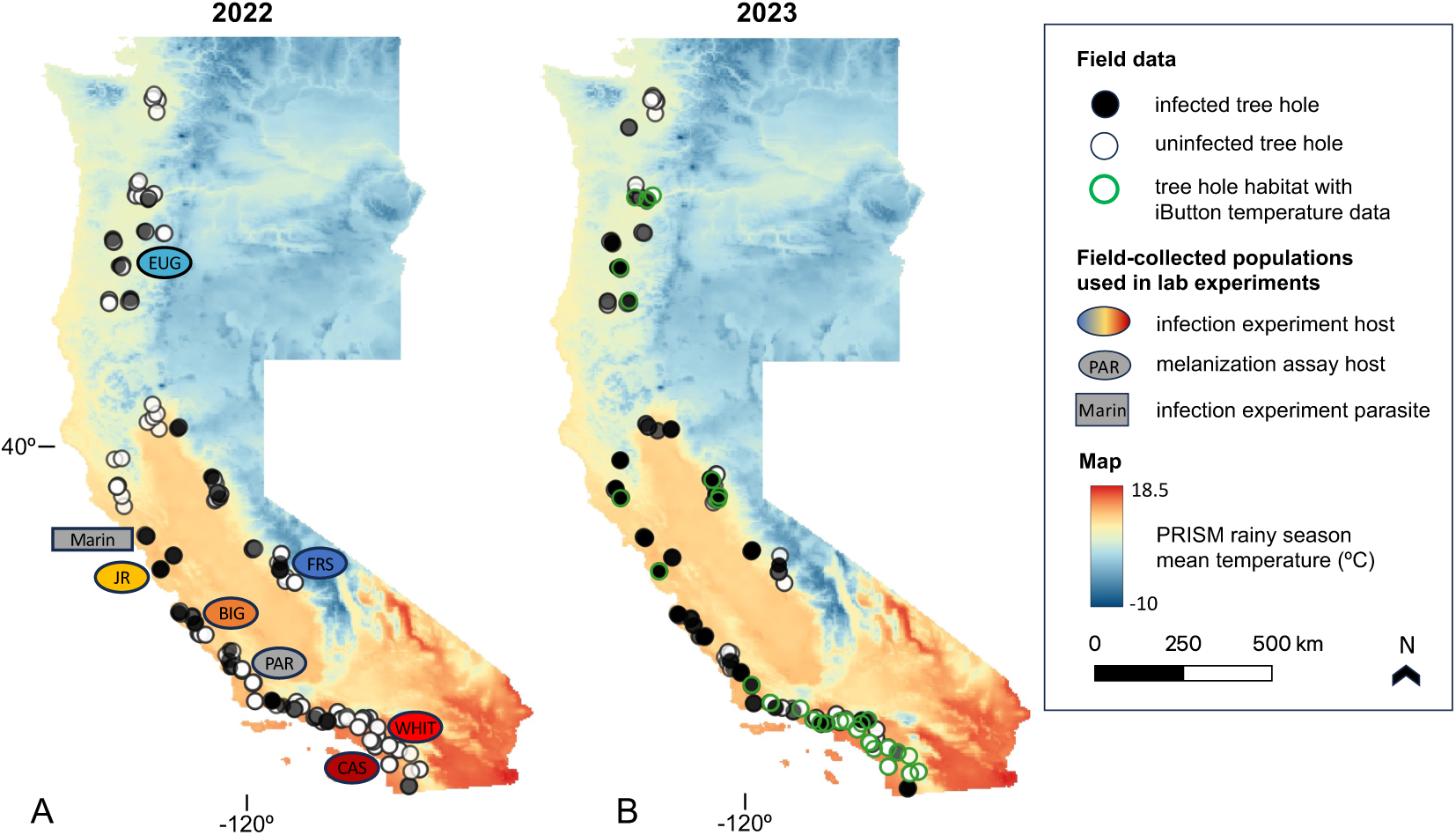
Tree holes were surveyed, and experimental host and parasite populations reared, from across a large geographic temperature gradient on the western coast of the United States during the (A) 2022 and (B) 2023 rainy seasons. The mapped color gradient shows mean temperature during the rainy season in Washington, Oregon, and California, USA, from the 4 km PRISM climate dataset. Points show the locations of 507 total surveyed tree holes. *L. clarki* infection presence is indicated by black points, and absence by white points. In (A), colored ovals indicate the source locations of the six host populations used in the infection experiment (from north to south: EUG = Eugene, OR; JR = Woodside, CA; FRS = Shaver Lake, CA; BIG = Big Sur, CA; WHIT = Trabuco Canyon, CA; CAS = San Juan Capistrano, CA); the gray oval shows the host population used in the melanization response experiment (PAR = Paso Robles, CA); and the gray rectangle shows the source location of the ciliate parasite culture used in the infection experiment (Marin = Novato, CA). In (B), green outlines indicate tree holes in which iButtons logged temperature data during the year between the two sampling visits.

For each sampling site, we used PRISM climate data to characterize long-term mean temperature conditions in the November – April rainy season when mosquito larvae and parasites are active in tree holes (PRISM Climate Group, 2021; Washburn & Anderson, 1986; Luković et al., 2021; Praskievicz & Chang, 2009). We chose the PRISM dataset for its long-term coverage of the entire study region. To assess the match between the remotely sensed PRISM data and the conditions in surveyed tree hole habitats, we placed iButton temperature loggers (Maxim Integrated, San Jose, CA, USA) in 84 arbitrarily selected tree holes across the survey area during the 2022 field season. These iButtons were retrieved the next year, and temperature logs were compared to PRISM estimates for the same locations during the intervening time period and found to be strongly correlated (R^2^ mean = 0.94, SD = 0.03). Finally, we accessed NOAA climate reports that compared regional temperatures and precipitation in each field season to long-term averages (NOAA National Centers for Environmental Information, 2022; 2023). Additional details on the field survey methods are provided in Appendix S1.

### Lab experiments

#### 1. Temperature-dependent infection experiment

In September 2022, we measured the effects of temperature on the host – parasite interaction with an experiment that used the F1 offspring of six host populations reared in the lab from larvae collected in the 2022 field survey: two populations each from warm (14.8 – 15.1°C rainy season mean temperature; San Juan Capistrano and Trabuco Canyon, CA), intermediate (10.8 – 11.8°C; Big Sur and Woodside, CA), and cool (5.3 – 6.5°C; Shaver Lake, CA and Eugene, OR) climates (Figure 1, Appendix S2: Table S1). We exposed each host population to the same allopatric parasite culture isolated from an infected larva collected in Marin County, CA (10.6°C) (Figure 1), so that any variation observed in the temperature dependence of infection would be due to the host population. Mosquito rearing and ciliate culturing occurred at 21.5°C following Ismail et al. (2023), Couper et al. (2024), and Lyberger et al. (2024).

For the experiment, mosquito eggs were hatched and day-old larvae were incubated in groups of five with 7 mL parasite culture diluted to a standardized density of 35 ciliates per 100 uL at six constant temperatures: 7°C, 11°C, 14°C, 18°C, 23°C, and 28°C (N = 6 microcosms per host population per temperature, except for Big Sur, where N = 4 due to lower egg hatching, totaling 34 microcosms per temperature, and 204 microcosms overall). We monitored the proportions of hosts with infections and parasite melanization spots, the survival of infected and uninfected hosts, and *L. clarki* cell density per 100 uL liquid up to the first pupation at each temperature. Because *Ae. sierrensis* development speed is thermally sensitive, we set monitoring frequencies that ensured at least four checkpoints spanning the host’s four larval instars at each temperature; these ranged from every two days at 28°C to every seven days at 7°C and 11°C (Appendix S2: Table S2).

#### 2. Host and parasite thermal responses in isolation

To compare the thermal performance of mosquito larvae and ciliates to that of parasitism, we incubated mosquito-only and ciliate-only microcosms alongside the infection experiment, following the same design as above. The mosquito-only microcosms were monitored on the same timeline as the infection experiment. For the ciliate-only microcosms (N = 6 wells per temperature, 36 total microcosms), we measured the population growth rate by inoculating wells with culture diluted to 4 cells per 100 uL and then counting the cells per 300 uL daily for four days of positive growth.

To assess the temperature sensitivity of the host melanization process (removing the temperature sensitivity of parasite attack rates), we performed a mechanical melanization assay on fourth instar larvae from one population at 4, 8, 12, 16, 20, 24, and 28°C (N = 6 wells of 5 mosquitoes per temperature with no ciliates, for 42 total microcosms). Additional details are provided in Appendix S1.

### Statistical analyses

#### 1. Field survey

To investigate the nonlinear temperature dependence of parasitism, we tested for a quadratic relationship between temperature and the presence of *L. clarki* infections in the surveyed tree holes using a binomial generalized linear mixed effects model, with mean temperature, mean temperature squared (to account for hypothesized nonlinear effects of temperature), and year as fixed effects, and tree hole ID as a random effect. We used the glmer function in the R package “lme4” with scaled and centered environmental variables, and assessed model fit with the R package “DHARMa” (Bates et al., 2003; Hartig, 2017). Because interactions between survey year and the temperature variables were not significant, we estimated the overall mean temperature where infection was most likely to occur with a model that did not include year. We additionally used general linearized models fit individually to the data from each survey year to describe the temperatures where infection occurrence peaked within each field season. To evaluate the relationship between within-tree hole habitat temperatures logged by iButtons and the PRISM air temperature dataset used in the model, we tested for correlations, calculated the mean differences between daily mean temperature time series recorded by iButtons and extracted from PRISM, and used linear regression with the mean temperature recorded within a tree hole as the dependent variable and the mean PRISM temperature estimate for that location as the independent variable. Because there were only two field seasons, the NOAA annual climate reports for each field season were not included in statistical analyses, but used only to qualitatively compare the two years.

#### 2. Lab experiments

To quantify the thermal sensitivity of the host – parasite interaction measured from the laboratory experiment, we used a Bayesian approach to fit TPCs for the proportion of infected hosts, the proportion of hosts with melanization spots, and the proportion of hosts killed by infections. The total number of infections observed in each microcosm over the course of the experiment depended on both initial parasite attack success and subsequent rounds of parasite transmission following infected host deaths that released new parasites as killed mosquitoes decomposed. We fit separate TPCs for both initial infection—the period up to the first deaths of infected hosts at each temperature, when all observed infections resulted from the first round of host – parasite interactions (covering checkpoints 1-3, approximately corresponding to larval instars 1-3)—and total infection rates over the entire experiment (including all host-parasite interaction cycles for all four larval instars); we used the same procedure to fit initial and total TPCs for the proportions of hosts melanizing attacking parasites. We additionally fit TPCs for free-living *L. clarki* population growth rate and for the host melanization immune response induced by mechanical penetration of the cuticle. We planned to fit TPCs for host survival in the absence of parasitism, but >99% of larvae incubated without parasites survived across all temperatures. Because initial parasite success is likely to determine whether there is a non-zero infection probability over the rest of the season (Washburn et al., 1991), this is the laboratory-based TPC that we expect to most closely align with the field-observed response of infection occurrence to temperature.

Unimodal traits were fit with a flexTPC functional form (Equation 1) which can describe unimodal TPCs of any skewness (Cruz-Loya et al., *preprint*). Using this flexible function to fit TPCs of different shapes across our traits of interest allowed for ease of comparison of thermal responses among traits and populations.

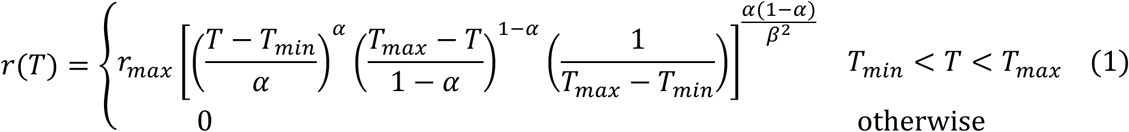

In this equation, 𝑇*_min_* and 𝑇*_max_* correspond to the minimum and maximum temperature, respectively, 𝑟*_max_* to the maximum trait value, and 𝛼 and 𝛽 are parameters that determine the curve skewness and rate of decrease from the maximum trait value, respectively (Appendix S1). Host melanization increased monotonically with temperature, and was fit to a 4-parameter shifted logistic model (Equation 2, see Appendix S1 for derivation).

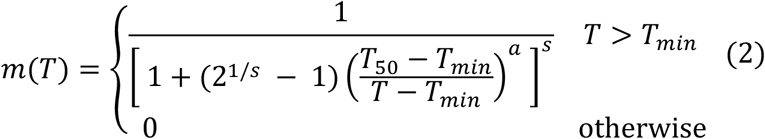

Here, 𝑇*_min_* is the minimum temperature, 𝑇_50_ is the temperature at which 50% of the population is melanized, and 𝑎 and 𝑠 are constants that allow for different rates of change of melanization near 𝑇_50_ and for the response to be asymmetric, which was necessary to fit the data.

We used a binomial likelihood for all traits that were measured as proportions. For the free-living parasite population growth rate TPC, we used a Gamma likelihood with population growth rates obtained from the raw time-course data with the R package “growthrates” (Petzoldt, 2022). Optimal temperatures were calculated analytically using the posterior samples of the model parameters (Appendix S1). Details on model fitting and diagnostics are provided in Appendix S1.

To assess population variation in lab-measured thermal performance among host populations and the responses measured in the infection experiment, we compared 95% credible intervals for the thermal optimum (T_opt_), maximum (T_max_), and minimum (T_min_) and interpreted non-overlapping intervals as evidence of biologically significant variation. We additionally compared the magnitudes of the 95% credible intervals over TPC plots for each population. Since the credible intervals for T_opt_, T_min_, and T_max_ overlapped for all populations for all measured traits, we combined the data to generate species-representative TPCs, which we used for our main analyses and interpretations. Finally, we compared the parasitism TPC to the host and parasite trait TPCs measured in isolation.

We used R version 4.2.1 for all statistical analyses and for figure generation (R Core Team, 2022). See Appendix S1 for additional details and R packages used.

## Results

### Does infection occurrence vary with temperature across the species range?

Infection occurrence followed a hump-shaped pattern along the surveyed temperature gradient, peaking at 9.4°C mean rainy season temperature (Figure 2). Mean temperature^2^ and year were significant predictors of *L. clarki* infection presence (Table 1). Quadratic models fit individually for infection occurrence in each year peaked at 10.0°C in 2022 and 9.2°C in 2023, but the interactions between year and both temperature variables were not statistically significant, indicating that the temperature – infection relationship did not vary between years (Table 1). Geographically, infections were common in northern California and southern Oregon and rarer at the warm and cold ends of the surveyed temperature gradient in southern California and northern Washington, respectively (Figure 1).

**Figure 2.**
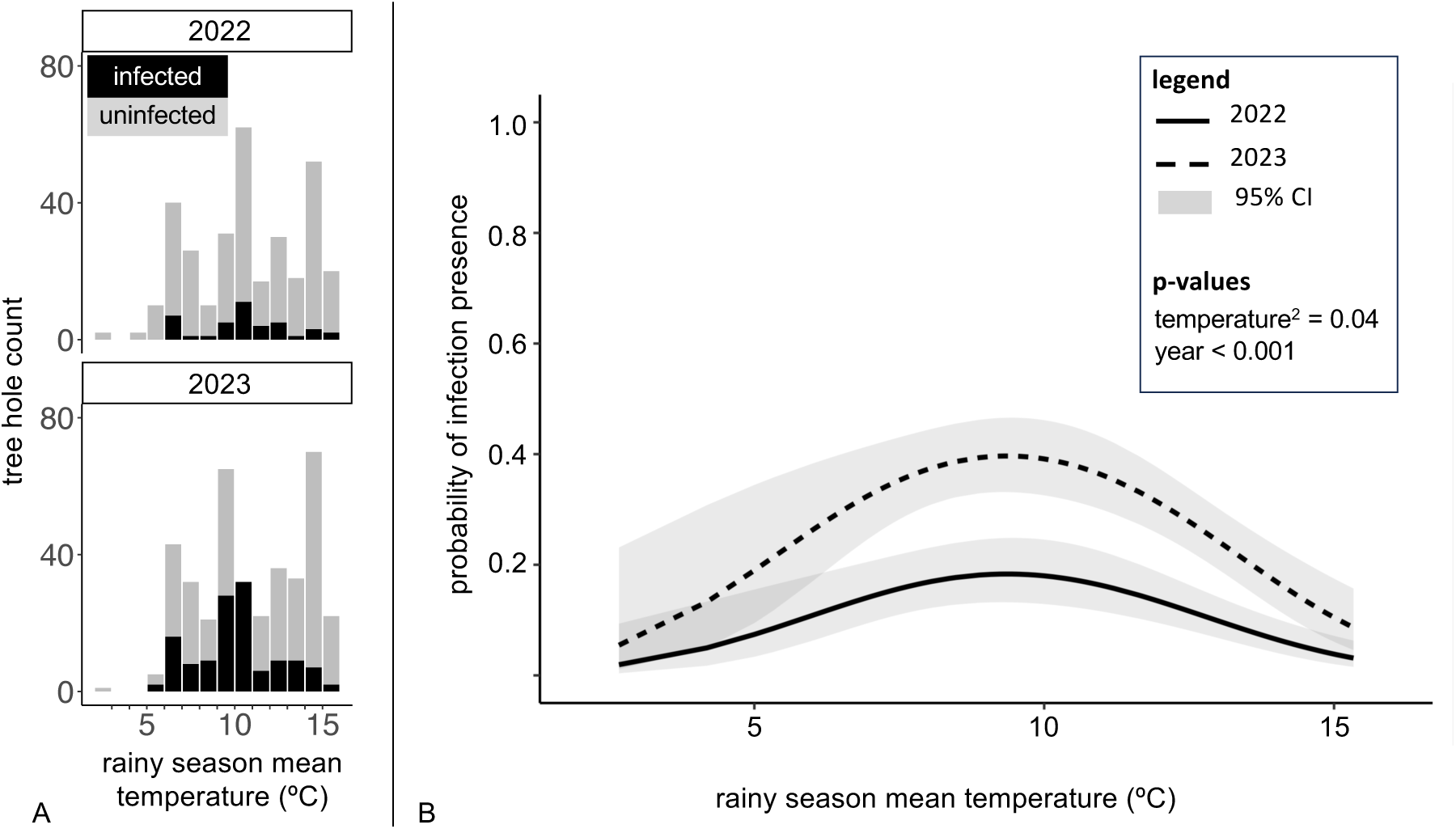
Infection probability followed a consistent hump-shaped relationship with mean temperature during the rainy season, and was higher overall in 2023 compared to 2022. **(A)** Counts of infected (black) and uninfected (gray) tree holes across the climate gradient in each survey year. **(B)** Generalized linear mixed-effects model predictions of infection probability by temperature and year are plotted across the surveyed climate gradient for each year of the field survey. The solid and dashed lines show the mean model predictions for 2022 and 2023, respectively, and the shading shows the 95% confidence intervals.

**Table 1.** Results of a generalized linear mixed-effects model with tree hole infection status as the dependent variable; mean rainy season temperature, mean rainy season temperature^2^, and survey year as fixed effects; and tree hole ID as a random effect.

| Predictor | Estimate | Standard error | Z-value | p-value |
| --- | --- | --- | --- | --- |
| *Mean rainy season temperature* | -0.22 | 0.22 | -1.01 | 3.13 x 10<sup>-1</sup> |
| *Mean rainy season temperature<sup>2</sup>* | -0.43 | 0.21 | -2.01 | **4.45 x 10<sup>-2</sup>** |
| *Survey year* | 1.31 | 0.34 | 3.90 | **9.5 x 10<sup>-5</sup>** |
| *Mean rainy season temperature \* survey year* | -0.45 | 0.26 | -1.72 | 8.5 x 10<sup>-2</sup> |
| *Mean annual temperature<sup>2</sup> \* survey year* | -0.13 | 0.25 | -0.52 | 6.34 x 10<sup>-1</sup> |

P-values < 0.05 are displayed in bold to highlight statistical significance.

Of the two sampling years, 2023 was cooler and wetter and had a higher percentage of infected tree holes (29.7%, N = 431) compared to 2022 (12.5%, N = 320) (Figure 2) (NOAA National Centers for Environmental Information, 2022; 2023). Mean tree hole habitat temperatures logged by iButtons (N = 32 successfully retrieved) were strongly correlated with and were not systematically warmer or colder than mean PRISM air temperature estimates (linear regression intercept = -0.41, SE = 1.76, t-value = -0.24, p-value = 0.82; slope = 0.95, SE = 0.1, t-value = 9.11, p-value = 3.86 x 10^-10^) (mean temperature difference = 1.3°C, SD = 2.6°C) (Appendix S2: Figure S1). Additional details are provided in Appendix S2.

### Does the thermal response of the host – parasite interaction depend on the host population?

We found similar thermal optima, minima and maxima across the six host populations in the infection experiment within each of the three measured interaction outcomes of infection, melanization of parasites, and mortality (Figure 3; Appendix S2: Figures S2-S11; Tables S3-S7). Although the mean estimated T_opt_, T_min_, and T_max_ varied slightly among populations for each trait, all 95% credible intervals across populations for each outcome x parameter combination overlapped. Therefore, we combined the data for all host populations to estimate broadly representative TPCs for the host – parasite interaction, with reduced uncertainty compared to the population-specific TPCs (Figure 4; Appendix S2: Figure S12; Table S8). The only trait in which we observed clear intraspecific variation was in mortality around 10°C, which was higher in one intermediate climate population (Big Sur (BIG)) compared to the other (Woodside (JR)) and to one cold climate population (Eugene (EUG)) (Figure S13).

**Figure 3.**
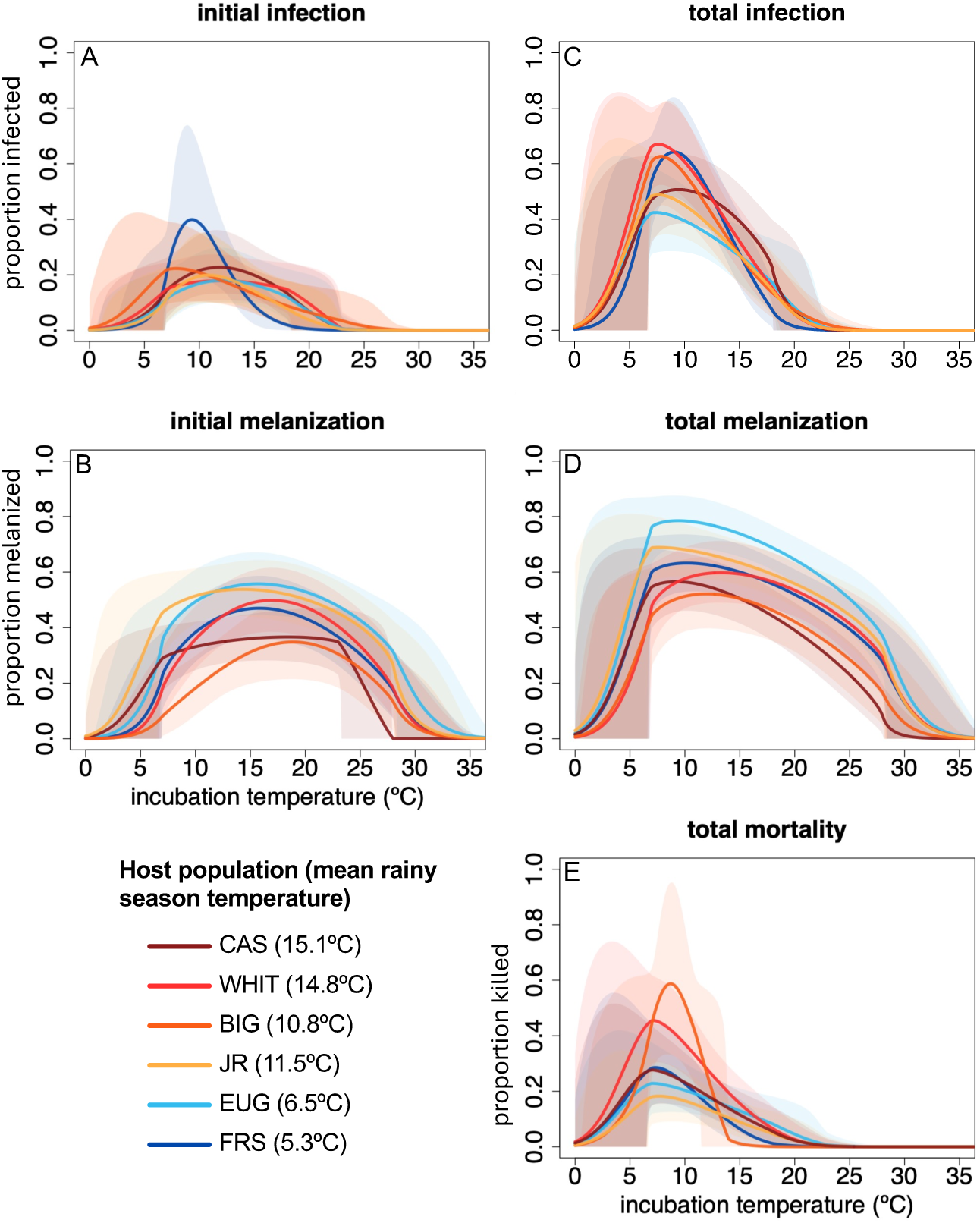
Thermal responses of host – parasite interaction outcomes were similar among host populations from warm, intermediate, and cold source climates. For each outcome measured in the infection experiment, mean estimated TPCs for each of six host populations are shown (lines) with 95% credible intervals (shading). The left column shows **(A)** initial infection and **(B)** parasite melanization responses up to the third experiment checkpoint, when the first deaths of infected larvae released additional parasites into the environment. The right column shows the **(C)** total infection, **(D)** parasite melanization, and **(E)** parasite-induced mortality rates across temperatures at the end of the experiment. Mortality rates were significantly higher near 10 °C for population BIG compared to populations JR and EUG. The TPCs shown here are presented individually, with the experiment data, in Appendix S2 (Figures S2, S4, S6, S8, S10).

**Figure 4.**
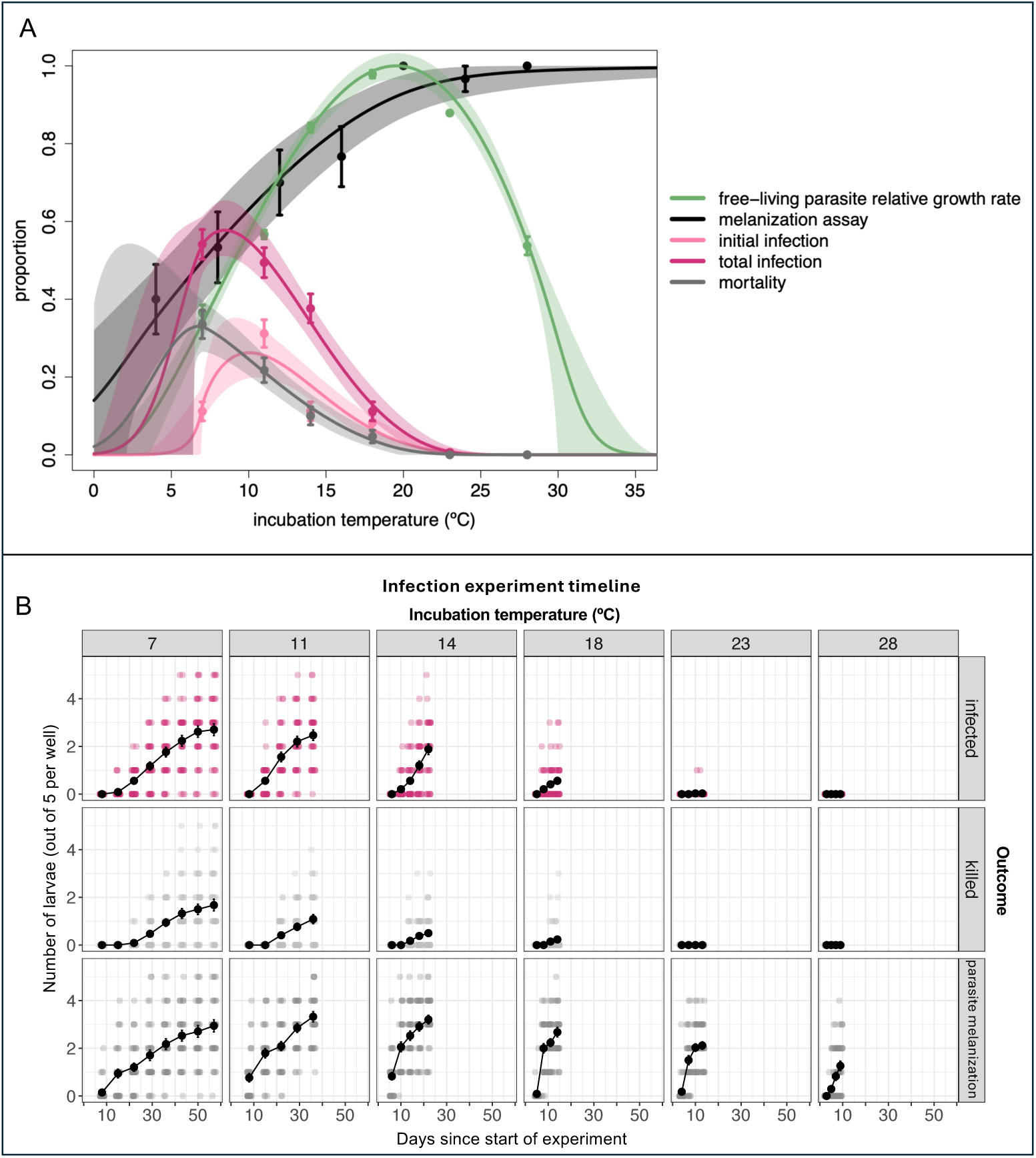
Experimental infections were most likely to occur at cold incubation temperatures where the host melanization response and development were suppressed. **(A)** Points show mean values, error bars show +/- 1 SE, lines show mean estimated TPCs, and shading shows 95% credible intervals for key host – parasitism interaction outcomes and processes: free-living parasite population growth (green); the host mechanically stimulated melanization immune response (black); the initial infection rate (light pink), calculated up to the first fatality at each temperature; the total infection rate (dark pink); and total mortality (gray) calculated up to the first pupation at each temperature. The infection data shown include all host populations in the infection experiment. **(B)** Infection (top; dark pink), parasite-induced mortality (middle; light gray), and parasite melanization (bottom; dark gray) counts are plotted over time for each temperature used in the infection experiment. Colored points show data from individual microcosms, and black points and lines show mean trajectories +/- 1 SE. For each temperature, the final experiment checkpoint was at the time of the first observed pupation.

### How do different components of the host – parasite interaction relate to the overall thermal sensitivity of parasitism?

Although free-living parasite population growth was maximized at warm temperatures, infection and parasite-induced mortality peaked at significantly colder temperatures, corresponding to conditions that suppressed host defense mechanisms, namely the melanization immune response and larval development speed (Figure 4). Specifically, the experimentally measured initial infection rate peaked at 10°C (95% CI: 8.7 °C, 11.3 °C), and the total infection rate peaked at 8.4°C (95% CI: 6.8 °C, 9.4 °C), much lower temperatures than the 19.5°C (95% CI: 18.9 °C, 20.2 °C) optimum for free-living parasite population growth, but near the field-estimated 10.0°C and 9.2°C peaks in 2022 and 2023, respectively (Figure 4A; Appendix S2: Figure S12; Tables S8-9). The mechanically assayed host melanization immune response increased with temperature up to 20°C, where it plateaued (Figure 4A; Appendix S2: Table S10). At warmer temperatures, attacking parasites faced not only a stronger host immune response, but also shorter time windows in which to infect hosts without being shed with the cuticle during larval molting and pupation or eaten by later larval instars. The time to the first observed pupation shortened as the temperature increased, occurring after 9 days at 28°C compared to 57 days at 7°C (Figure 4B), consistent with previously measured *Ae. sierrensis* larval development rate TPCs that peaked at 27.7°C (Couper et al., 2024). Consequently, rapid pupation interrupted upward trajectories of infection and mortality at intermediate temperatures, contributing to the decline in parasitism above 11°C, and to the lower T_opt_ of total compared to initial infection. At colder temperatures, additional opportunities for parasitism were also generated as infected host deaths beginning in the third week of the experiment maintained higher *L. clarki* densities throughout extended host developmental periods (Figure S14). In comparison, at the warmest temperature with infections (18°C), the contributions of infected host deaths to subsequent transmission were limited by their rarity and proximity to the onset of pupation.

## Discussion

In this study, we used complementary field observations and lab experiments to understand how temperature affects a host-parasite interaction across a large, climatically variable species range. We found that infection peaked at sites with intermediate temperatures (9.4°C) in the field across two sampling years and was higher in a colder, wetter year, aligning with the unimodal temperature-dependence of infection measured in the lab, which peaked at 10°C. This reflects the interacting thermal biology of multiple mosquito and ciliate traits, which together led to an optimal temperature for parasitism (8-10°C for total versus initial infection) that was colder than the optimal temperature for free-living parasite performance (19.5°C). Infection rates of mosquito populations from different climates had similar thermal responses in the lab, and a single TPC described the infection process across the entire surveyed area. These findings suggest that for systems with strong temperature dependence, TPCs measured under simplified lab conditions can capture patterns of parasitism across large geographic temperature gradients in complex natural settings, by quantifying the underlying thermal biology mechanisms.

### Lab experiments capture nonlinear parasitism thermal responses in the field

The results add to evidence that temperature nonlinearly affects parasitism not only in the lab, but also in nature, making laboratory and field studies that are designed to capture nonlinear temperature effects valuable tools for understanding disease risk (Dudney et al., 2021; Kirk et al., 2024; Shocket, Ryan et al., 2018; Shocket, Strauss et al., 2018). However, despite the similarities between the field observations and the lab-measured parasitism TPC, we cannot rule out the influence of other factors. For example, while the role of precipitation in this system has not been assessed, the colder survey year with higher infection prevalence was also wetter, suggesting that precipitation patterns that determined when and for how long tree holes were water-filled may have also affected the potential for parasitism to occur. Beyond mean temperature differences, heatwaves or cold snaps occurring during key points in the host – parasite interaction may have also contributed to interannual variation in infection presence (Claar & Wood, 2020; Altman et al., 2016). These limitations should be addressed with field experiments that manipulate temperatures in natural tree holes at different latitudes or elevations; lab experiments that incorporate additional environmental complexity; and long term repeated yearly sampling of the same tree holes.

### Intraspecific variation

Very few studies investigate the impact of local thermal adaptation on species distributions and host – parasite interactions, a gap often noted in the literature (Sternberg & Thomas, 2014; Urban et al., 2016; Couper et al., 2021; Aleuy et al., 2023). In addressing this gap, we did not find evidence of local temperature adaptation affecting infection patterns across temperatures in the lab or the field. While our laboratory experiment specifically assessed host local adaptation, the invariant temperature dependence of infection we observed is consistent with the similar temperature-infection relationship in the field, and with our previous work comparing sympatric and allopatric host – parasite interactions. Specifically, for nine *Ae. sierrensis* and *L. clarki* populations with free-living parasite T_opt_ ranging from 16.5°C to 23.5°C, infection rates measured at three temperatures peaked at the intermediate temperature of 12.5°C, substantially cooler than the free-living growth optima (Lyberger et al., 2024). Cumulatively, these findings suggest that although both *Ae. sierrensis* pupal development and *L. clarki* free-living population growth rates are locally adapted to temperature, this intraspecific variation is unlikely to directly affect host – parasite interactions (Ismail et al., 2023; Couper et al., 2024; Lyberger et al., 2024). Variability in the infection T_opt_ should be further tested through TPC experiments with additional host – parasite population pairs, but differences in factors such as host fitness and adaptation to parasites may be more important sources of intraspecific variation in parasite impacts. For example, in this experiment, the mosquito population with the lowest egg hatching rate, BIG, had a higher mortality rate near the infection T_opt_ compared to two other populations; and in previous experiments, individual host populations varied in their susceptibility to different parasite strains (Ismail et al., 2023; Lyberger et al., 2024)

Importantly, this is one of the only ectothermic host – parasite systems for which interaction thermal sensitivity has been assessed for populations from different climates, along with a stickleback – tapeworm system in which effects of temperature on parasitism were similar among four study populations collected from across a similar geographic temperature gradient (Franke et al., 2019). Notably, intraspecific variation in parasitism temperature sensitivity has been shown for levels of biological organization both above and below the population level that we assessed, for *Batrachochytrium dendrobatitis* infection both among different frog host species, and among salamander host color morphs within a single population (Cohen et al., 2017; Venesky et al., 2022). However, if population thermal sensitivities of parasitism are commonly consistent across species ranges, broadly applicable TPCs should be easily attainable for the many host – parasite systems that are amenable to laboratory studies, and field studies may also be able to capture nonlinear responses of parasitism to temperature.

### Mechanisms of parasitism temperature dependence

Our finding that parasitism peaked at temperatures that constrained host development and defense—rather than at the thermal optimum for parasite growth and attack—aligns with theory on the thermal biology of biotic interactions. Thermal asymmetry theory posits that parasitism peaks at temperatures where parasites outperform hosts, which may differ from optimal temperatures for the parasites themselves (Dell et al., 2014; Cohen et al., 2017). Our study supports the value of approaching this problem as the outcome of multiple interacting host and parasite traits with varying thermal responses (Paull et al., 2012; Shocket et al., 2018; Sun et al., 2023).

## Conclusions

Overall, our characterization of the temperature – infection relationship of the *Ae. sierrensis* host – *L. clarki* parasite interaction across settings demonstrates how controlled lab experiments can be paired with large-scale field observations to understand the complex thermal biology of parasitism. Climate warming is expected to cause geographic contractions, expansions, and shifts in host thermal refuges from disease (Altizer et al., 2013; Gsell et al., 2023). Our results suggest that parasite burdens on *Ae. sierrensis* may decline at southern latitudes and low elevations as thermally suitable conditions for parasitism shift northwards and upwards into areas where cold conditions currently limit *L. clarki* infections. Whether the distribution of parasitism ultimately tracks climate warming will depend on multiple factors, including parasite dispersal, the pace of thermal adaptation in host and parasite traits that allow local populations of each species to persist individually, and the effects of other environmental changes in conjunction with increasing mean temperatures, such as increasingly extreme precipitation patterns (Swain et al., 2018). Understanding how temperature affects parasitism in inherently complex and variable natural systems remains a pressing problem with broad implications for current and future disease risk management. Simple pairings of field surveys and lab experiments can offer straightforward means for approaching this challenge.

## Acknowledgments

Otto Seppälä, Jan Washburn, Nona Chiariello, and Angie Nakano played critical roles in establishing the study system. Our field sampling and mosquito rearing efforts were made possible by assistance from staff at vector control agencies for the California counties of San Mateo, Santa Clara, Marin-Sonoma, Solano, Alameda, Shasta, Placer, Santa Barbara, and Riverside; Jasper Ridge Biological Preserve ‘Ootchamin ‘Ooyakma; the UC Reserves at Emerson Oaks, Dawson Los Monos Canyon, and Hopland; and Andrew Rivera of Clarke Pest Control. Mordecai lab members helped with setting up the infection experiment. JEF was funded by the Bing-Mooney Fellowship and the ARCS Foundation. EAM and MCL were supported by the National Institutes of Health (R35GM133439 and R01AI168097). EAM was supported by the National Science Foundation (DEB-2011147 with Fogarty International Center), the National Institutes of Health (R01AI102918), the Stanford Woods Institute for the Environment, the Stanford Center for Innovation in Global Health, and the Stanford Doerr School of Sustainability. KL was funded by the NSF Postdoctoral Research Fellowships in Biology Program under Grant No. 2208947. LIC was funded by the NSF Postdoctoral Research Fellowship in Biology. MCL was supported by a Stanford PRISM Baker Fellowship.

