## Appendix S1 for "Nonlinear effects of temperature on mosquito parasite infection across a large geographic climate gradient" (from another version of this preprint: version 2, 2025-07-19)

Ecology

#### Appendix S1: Methods

### Field survey

Across the study region, we used satellite imagery, web searches, and guidance from local vector control agencies to identify accessible areas with tree cover that were likely to contain tree hole habitats, and then searched for and opportunistically collected samples from tree holes with *Ae. sierrensis* larvae present. The lowest and highest latitudes sampled were at 32.68°N and 47.76°N, respectively. The lowest elevations sampled were at sea level along the Pacific coast, and the highest elevations sampled were at 1370m on the western slopes of the Sierra Nevada and Cascade Mountains. In order to slow processes including mosquito development, infection development, and infection clearance, all samples were kept chilled from the point of collection until inspection for parasite infections at 10-40x magnification with a dissection microscope. For each sample, juvenile mosquitoes (larvae and pupae) were checked for infections until either infection was observed, the sample was exhausted, or 50 individuals had been inspected.

We used Google Earth Engine to extract long-term mean November to April rainy season temperature and precipitation data for the location of each sampled tree hole from the 4km spatial resolution PRISM climate dataset for 1991 – 2020, and separately for the 2022 and 2023 rainy seasons of the field survey (PRISM Climate Group 2021; Funk et al. 2015).

Ibutton temperature loggers (Maxim Integrated DS1921G, San Jose, CA) placed in tree holes were programmed to log data every four hours. Following iButton retrieval, within-tree hole daily mean temperature time series were compared to matching daily mean air temperature time series extracted from the 4km PRISM climate dataset using Google Earth Engine.

### Lab experiments

#### 1. Materials

Each experimental population originated from a single tree hole at least 20km distant from all other tree holes from which materials were used in experiments. The Marin County tree hole from which the experimental parasite culture was sourced was >80km apart from the source locations of all experimental mosquito populations. To ensure sufficient egg counts, genetic variation and an absence of *L. clarki* contamination in the mosquito eggs used in the experiment, we reared mosquitoes only from uninfected tree hole samples of >30 larvae.

#### 2. Experiments

All experimental microcosms were housed in six-well plates (Corning #3736). Two incubators were maintained at each experimental temperature, and the microcosms in each experimental group were split evenly between these. The plates within each incubator were shuffled twice weekly. For the infection experiment, each microcosm contained 5 mosquito larvae, 7 mL ciliate parasite culture diluted to a density of 35 cells per 100 uL, 1 mL hatching water, and

1 autoclaved barley seed. In the host-only experiments, 7 mL of autoclaved culturing media was used in place of parasite culture. In the parasite-only experiment, the 7 mL of culture were diluted to a low density of 4 cells per 100 uL. We used a dissection microscope at 10 – 40x magnification to monitor infections, melanization, and ciliate presence.

##### Melanization assay

We challenged the immune system of larvae by piercing the thoracic cuticle with a piece of 0.08-mm diameter wire cut at an angle to create a sharp point, avoiding internal organs. Larvae were then immediately returned to incubators and checked for puncture wound melanization after 24 hours. To maintain consistency, this procedure was performed by a single researcher. Due to limited egg stocks for the mosquito populations used in the main experiment, we used a population from Paso Robles, in central California (PAR; 10.9ºC) (Figure 1). We expect the temperature dependence of melanization in PAR to be representative of the species, as temperature effects were consistent across populations in our infection experiment (see below), and in a previous comparison of *Ae. sierrensis* larval trait TPCs across the study region (Couper et al. 2024).

### Data analysis

In addition to the R packages listed in the main text and below, we used "corrplot", "ggeffects", "ggforce", "mcmcplots", "MCMCvis", and "tidyverse" for data analysis and figure generation (Wei et al. 2017; Lüdecke 2018; Pedersen 2024; Wickham et al. 2019; Curtis, Goldin, and Evangelou 2018; Youngflesh 2018). We used QGIS to plot the maps shown in Figure 1 (QGIS Development Team 2024).

### Thermal performance curves

In this section, we describe the methodology used for fitting thermal performance curves. As described in the main text, we first fit TPCs for each population separately. Because the TPC fits were similar across all populations for each trait, we then combined the data across all populations to fit a single representative TPC for each trait. This second set of TPCs was used for our main analysis.

We fit models using Markov Chain Monte Carlo (MCMC) with the r2jags R package, an interface for the JAGS (Just Another Gibbs Sampler) program (Plummer 2003; Su et al. 2015). Four independent MCMC chains were run for 1,000,000 iterations, discarding the first 100,000 iterations as burn-in. The resulting MCMC chains were thinned, saving every eight iterations. Chain convergence was monitored both by visual inspection of trace plots and density plots of the individual chains and by ensuring the potential scale reduction factor

$\hat{R} < 1.01$ for all parameters. Posterior summaries for all TPC fits are provided in Appendix S2.

#### Unimodal traits

Traits that had a unimodal response to temperature (initial and final infection proportions, parasite melanization proportions, mortality and parasite growth rate) were modeled with a flexTPC functional form (Cruz-Loya, Mordecai, and Savage 2024)

$$r(T) = \begin{cases} r_{\max}\left[\left(\frac{T-T_{\min}}{\alpha}\right)^{\alpha}\left(\frac{T_{\max}-T}{1-\alpha}\right)^{1-\alpha}\left(\frac{1}{T_{\max}-T_{\min}}\right)\right]^{\frac{\alpha(1-\alpha)}{\beta^2}} & T_{\min} < T < T_{\max} \\ 0 & \text{otherwise} \end{cases} \tag{1}$$

which can describe thermal performance curves of left-skewed, symmetrical or right-skewed traits. This model is parametrized in terms of the minimum temperature $T_{\min}$, the maximum temperature $T_{\max}$, the maximum trait value $r_{\max}$, the curve skewness parameter $\alpha$ which determines the location of the thermal optimum, and the breadth parameter $\beta$ that determines the rate at which trait performance decreases from its maximum value.

##### Statistical model for proportions

Traits that are proportions (initial and final infection proportions, parasite melanization proportions and mortality) were modeled with a binomial likelihood:

$$n_i | T_i, N_i, \mathcal{P} \sim \text{Binomial}(p = r(T_i; \mathcal{P}), N = N_i)$$

where $N_i$ is the total number of individuals evaluated at temperature $T_i$, $n_i$ the number of individuals positive for the trait evaluated, and $r(T_i; \mathcal{P})$ is the flexTPC model (Equation 1) evaluated at temperature $T_i$ with parameters $\mathcal{P} = \{T_{\min}, T_{\max}, r_{\max}, \alpha, \beta\}$.

The following prior distributions were used for the flexTPC model parameters:

$$\begin{aligned} T_{\min} &\sim \text{Normal}(\mu = 5°\text{C}, \sigma = 2.5°\text{C}) \\ T_{\max} &\sim \text{Normal}(\mu = 25°\text{C}, \sigma = 5°\text{C}) \\ r_{\max} &\sim \text{Uniform}(0, 1) \\ \alpha &\sim \text{Uniform}(0, 1) \\ \beta &\sim \text{Gamma}(\mu = 0.3, \sigma = 0.3) \end{aligned}$$

The priors for $T_{\min}$ and $T_{\max}$ were chosen to make conservative assumptions on the possible temperature range for these parameters informed by our previous work on the thermal biology of *Ae. sierrensis* and *L. clarki*. Specifically, we assumed that $T_{\min}$ is *a priori* 95% likely to be between 0°C, where habitat freezing should prevent parasite attack, and 10ºC, below which we have previously observed low infection rates (Lyberger et al. 2024; Ismail et al. 2023). Similarly, we assumed a 95% prior probability of $T_{\max}$ falling between 15°C, above which we have previously observed declining infection rates, and 35°C, a temperature above the previously observed critical thermal limit of larval survival for *Ae. sierrensis* (Ismail et al. 2023; Lyberger et al. 2024; Couper et al. 2024). The prior for $r_{\max}$ is a noniformative choice that corresponds to the *a priori* assumption that the maximum value of the trait (which is a proportion) is equally likely to be in the interval $[0, 1]$.

FlexTPC has two additional parameters that determine the shape of the curve. Parameter $\alpha$ determines the location of the thermal optimum through the equation $T_{\text{opt}} = \alpha T_{\max} + (1-\alpha)T_{\min}$, where, for example, $\alpha = 0$ corresponds to $T_{\text{opt}} = T_{\min}$, $\alpha = 1$ to $T_{\text{opt}} = T_{\max}$ and $\alpha = \frac{1}{2}$ to a symmetric TPC where $T_{\text{opt}} = \frac{T_{\min}+T_{\max}}{2}$. The uniform prior on $\alpha$ is also an uninformative choice that makes the assumption that $T_{\text{opt}}$ is *a priori* equally likely to be anywhere in the interval $[T_{\min}, T_{\max}]$. Parameter $\beta > 0$ determines whether the TPC is narrow (i.e., with performance rapidly decreasing near the optimum, which corresponds to small $\beta$) or broad (i.e., with performance that slowly decreases near the optimum, corresponding to large $\beta$). The prior chosen for this parameter is weakly informative, preferring values of $\beta$ that correspond to curves with a similar broadness to those of other common TPC models (such as quadratic or Briere1, which correspond to $\beta$ of around 0.2 to 0.4), but allowing other values in case they are necessary for describing the data.

##### Statistical model for *Lambornella* growth rates

Parasite growth rates were modeled with a Gamma likelihood

$$y_i | T_i, \sigma, \mathcal{P}_i \sim \text{Gamma}(\mu = r(T_i; \mathcal{P}_i), \sigma = \sigma)$$

to account for growth rates being nonnegative. The priors used for the flexTPC model were as follows:

$$\begin{aligned}
T_{\min} &\sim \text{Normal}(\mu = 5°\text{C}, \sigma = 2.5°\text{C}) \\
T_{\max} &\sim \text{Normal}(\mu = 25°\text{C}, \sigma = 5°\text{C}) \\
r_{\max} &\sim \text{Exponential}(\mu = 10) \\
\alpha &\sim \text{Uniform}(0, 1) \\
\beta &\sim \text{Gamma}(\mu = 0.3, \sigma = 0.3)
\end{aligned}$$

The priors for $T_{\min}$, $T_{\max}$, $\alpha$ and $\beta$ are identical to those described previously for traits that are proportions. A different prior was used for the maximum trait value $r_{\max}$ as, unlike proportions, growth rates can be greater than one. A diffuse exponential prior was chosen with the goal of it having little to no influence in the parameter estimates.

In addition to the parameters of the TPC model, the use of a Gamma likelihood requires an additional prior for the standard deviation of the observations:

$$\sigma \sim \text{Uniform}(0, 1)$$

This is an uninformative choice that is intended to not have a strong effect in parameter inference.

### Monotonic traits

While most traits exhibited a unimodal relationship with temperature, we found a monotonically increasing relationship between temperature and host melanization. This trait was modeled with a four-parameter shifted logistic model (see mathematical derivation below)

$$m(T) = \begin{cases} \frac{1}{\left[1+\left(2^{\frac{1}{s}}-1\right)\left(\frac{T_{50}-T_{\min}}{T-T_{\min}}\right)^{a}\right]^{s}} & T \geq T_{\min} \\ 0 & \text{otherwise} \end{cases} \tag{2}$$

where $T_{\min}$ is the minimum temperature for melanization, $T_{50}$ the temperature at which half the population is melanized and $a, s > 0$ are arbitrary constants that allow for the response near $T_{50}$ to vary at different rates and possibly be asymmetric. The shifted logistic model for this trait was chosen after initial attempts to use a Hill-function based model did not provide a satisfactory fit to the data due to the asymmetry in the observed melanization response.

#### Statistical model

We modeled host melanization with a binomial likelihood:

$$n_i | T_i, N_i, \mathcal{P} \sim \text{Binomial}(p = m(T_i; \mathcal{P}), N = N_i)$$

where $N_i$ is the total number of individuals evaluated at temperature $T_i$, $n_i$ the number of individuals positive for melanization, and $m(T_i; \mathcal{P})$ is the shifted logistic model evaluated at temperature $T_i$ with parameters $\mathcal{P} = \{T_{\min}, T_{50}, a, s\}$.

The following prior distributions were used for the TPC model:

$$\begin{aligned} T_{\min} &\sim \text{Normal}(\mu = 0°\text{C}, \sigma = 5°\text{C}) \\ T_{50} &\sim \text{Normal}(\mu = 20°\text{C}, \sigma = 10°\text{C}) \\ \log a &\sim \text{Normal}(\mu = 0, \sigma = 1) \\ \log s &\sim \text{Normal}(\mu = 0, \sigma = 1) \end{aligned}$$

The prior for $T_{\min}$ corresponds to the assumption that, *a priori*, the minimum temperature for melanization is 95% likely to be in the interval $[-10°\text{C}, 10°\text{C}]$. This includes a wider range of negative temperatures compared to other traits. This was based on a lack of strong evidence that melanization rates in insects decline to zero at 0°C, given that *Aedes* mosquitoes exhibit chill coma adaptations to cold climates that allow larval survival at subfreezing temperatures (Jass et al. 2019); that immune activity occurred at the lowest temperature we measured, and has been observed for temperatures as low as 0.5°C in the cricket *Gryllus veletis* (Ferguson, Heinrichs, and Sinclair 2016); and that the thermal responses of the biochemical processes that eventually kill mosquito larvae at freezing temperature may differ from the processes that affect melanization. The prior for $T_{50}$ is weakly informative but very biologically conservative, assuming that the temperature at which half of the population exhibits melanization is *a priori* 95% likely to be in the interval $[0°\mathrm{C}, 40°\mathrm{C}]$. The priors on $\log a$ and $\log s$ are also weakly informative, corresponding to the assumption that, *a priori*, $a$ and $s$ are 95% likely to be in the interval $[10^{-2}, 10^{2}]$.

### Mathematical derivation of shifted logistic model

Monotonic relationships between a response $r$ and the concentration of a chemical $x$ are often modeled by the Hill equation

$$r(x) = r_{\min} + \frac{r_{\max} - r_{\min}}{1 + \left(\frac{c}{x}\right)^{a}}$$

where $r_{\min}$ and $r_{\max}$ are the minimum and maximum value of the response, $c$ is the concentration at which 50% of the response is attained and $a$ is an arbitrary constant. This model assumes that the minimum value is attained when $x = 0$ (i.e. when the chemical is absent) and that it approaches an asymptote at $r_{\max}$ as $x \to \infty$.

The Hill equation is symmetrical around its midpoint, with the rate of change of the response being approximately the same above and below the midpoint. To model monotonic relationships that can be asymmetric, the Hill model can be extended by adding a parameter $s$, which yields the five-parameter logistic model (Prentice 1976)

$$r(x) = r_{\min} + \frac{r_{\max} - r_{\min}}{\left[1 + \left(\frac{c}{x}\right)^{a}\right]^{s}}$$

where, as before, $r_{\min}$ and $r_{\max}$ are the minimum and maximum value of the response and $a, s$ are arbitrary constants. This model includes the Hill model as a special case when $s = 1$. The interpretation of $c$ changes in the five-parameter logistic model, as it is no longer the concentration where 50% of the response is attained (this is only true when $s = 1$). Instead, it becomes another arbitrary constant.

In this work, we modify this five parameter logistic model as follows to make it suitable for describing a monotonically increasing temperature performance curve:

- We replace concentration $x$ with $T - T_{\min}$, where $T$ represents temperature and $T_{\min}$ the minimum temperature at which trait performance changes with temperature.

- We reparametrize the model to replace $c$ with $T_{50}$, the temperature at which 50% of the maximum response is attained.

This yields the six-parameter model

$$r(T) = \begin{cases} r_{\min} + \frac{r_{\max} - r_{\min}}{\left[1+\left(2^{\frac{1}{s}}-1\right)\left(\frac{T_{50}-T_{\min}}{T-T_{\min}}\right)^{a}\right]^{s}} & T \geq T_{\min} \\ r_{\min} & \text{otherwise} \end{cases}$$

To keep the usual definition of minimum temperature, we further assume that $r(T)$ will be zero at $T_{\min}$ (i.e. that $r_{\min} = 0$). Also, we remove another parameter by assuming that the proportion of melanized hosts approaches 1 at high remperatures (i.e., $r_{\max} = 1$). This yields the four-parameter shifted logistic model that was used to fit the host melanization temperature curves (Equation 2).

[11] Pedersen, Thomas Lin. 2024. "Ggforce: Accelerating 'Ggplot2.'" R Package Version 0.4.
