## Appendix S2 for "Nonlinear effects of temperature on mosquito parasite infection across a large geographic climate gradient" (from another version of this preprint: version 2, 2025-07-19)

*Ecology*

**Appendix S2: Results, Figures, and Tables**

Nonlinear effects of temperature on mosquito parasite infection across a large geographic climate gradient

Johannah E. Farner, Kelsey P. Lyberger, Lisa I. Couper, Mauricio Cruz-Loya, Erin A. Mordecai

### Results

*Field survey*

We located and sampled a total of 507 tree holes with *Ae. sierrensis* present over the two years of the field survey (Figure 1). Totals of 320 and 431 tree holes containing *Ae. sierrensis* were sampled in 2022 and 2023, respectively, 244 of which were sampled in both years. We observed *L. clarki* infections in 12.5% of sampled tree holes in 2022 (n = 40 infected tree holes out of 320), and 29.7% of sampled tree holes in 2023 (n = 128 infected tree holes out of 431) (Figures 1-2). Eighteen of the tree holes that were sampled twice were positive for infections in both years of the survey (56.3% of all tree holes positive for infection at least once). Long-term (1991 – 2020) mean rainy season temperatures at the surveyed sites ranged from 2.7ºC to 15.3 ºC (mean = 10.7ºC, SD = 2.8ºC) (PRISM Climate Group and Oregon State University 2021). NOAA reported that rainy season conditions in 2022 were warmer and drier than average in California and Oregon; in 2023, the entire region was colder than average, and California was wetter than average (NOAA National Centers for Environmental Information 2022; 2023). For the 2022 rainy season, PRISM estimated mean temperatures for sampled tree holes ranged from 3.6ºC to 15.9ºC (mean = 11.3ºC, SD = 3.1ºC), and cumulative precipitation estimates ranged from 158mm to 1707mm (mean = 467mm, SD = 247mm) (PRISM Climate Group and Oregon State University 2021). For the 2023 field season, rainy season mean temperature estimates ranged from -0.5ºC to 13.2ºC (mean = 9.3ºC, SD = 2.5ºC), and cumulative precipitation estimates ranged from 393mm to 2117mm (mean = 1038mm, SD = 328mm) (PRISM Climate Group and Oregon State University 2021).

### Figures

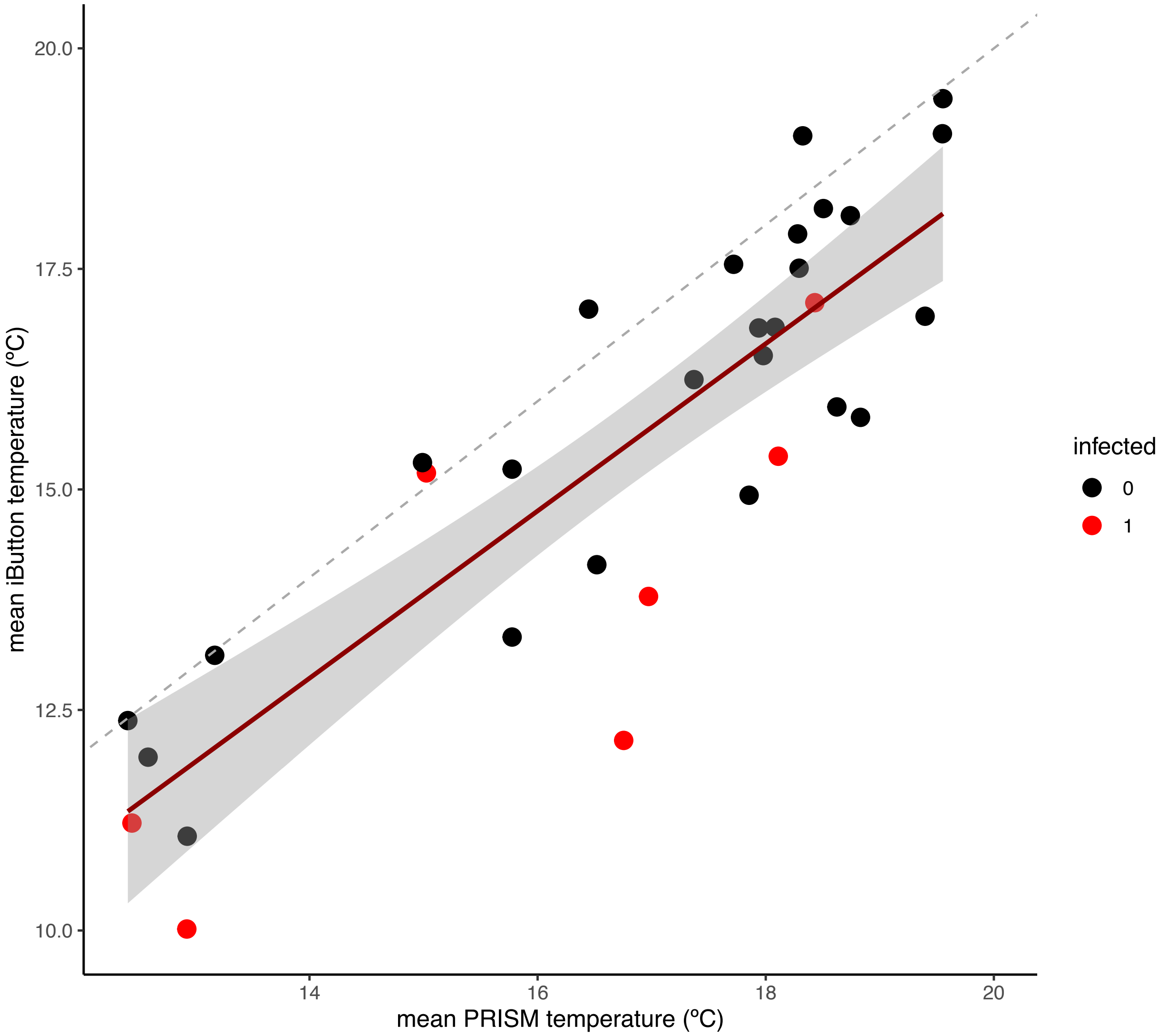


**Figure S1. The mean daily temperatures recorded by iButtons placed within tree holes were strongly correlated with, but usually cooler than, the corresponding PRISM mean daily temperatures for the same time period.** Each point represents a sampled tree hole. Red points indicate tree holes with *Lambornella clarki* infections present in at least one year of the survey. The dashed grey line shows the line of equality, and the dark red line was fit with linear regression (estimated slope = 0.95, p-value = $3.86 \times 10^{-10}$; estimated intercept = -0.41, p-value = 0.82) (grey shading = +/- 1 SE).

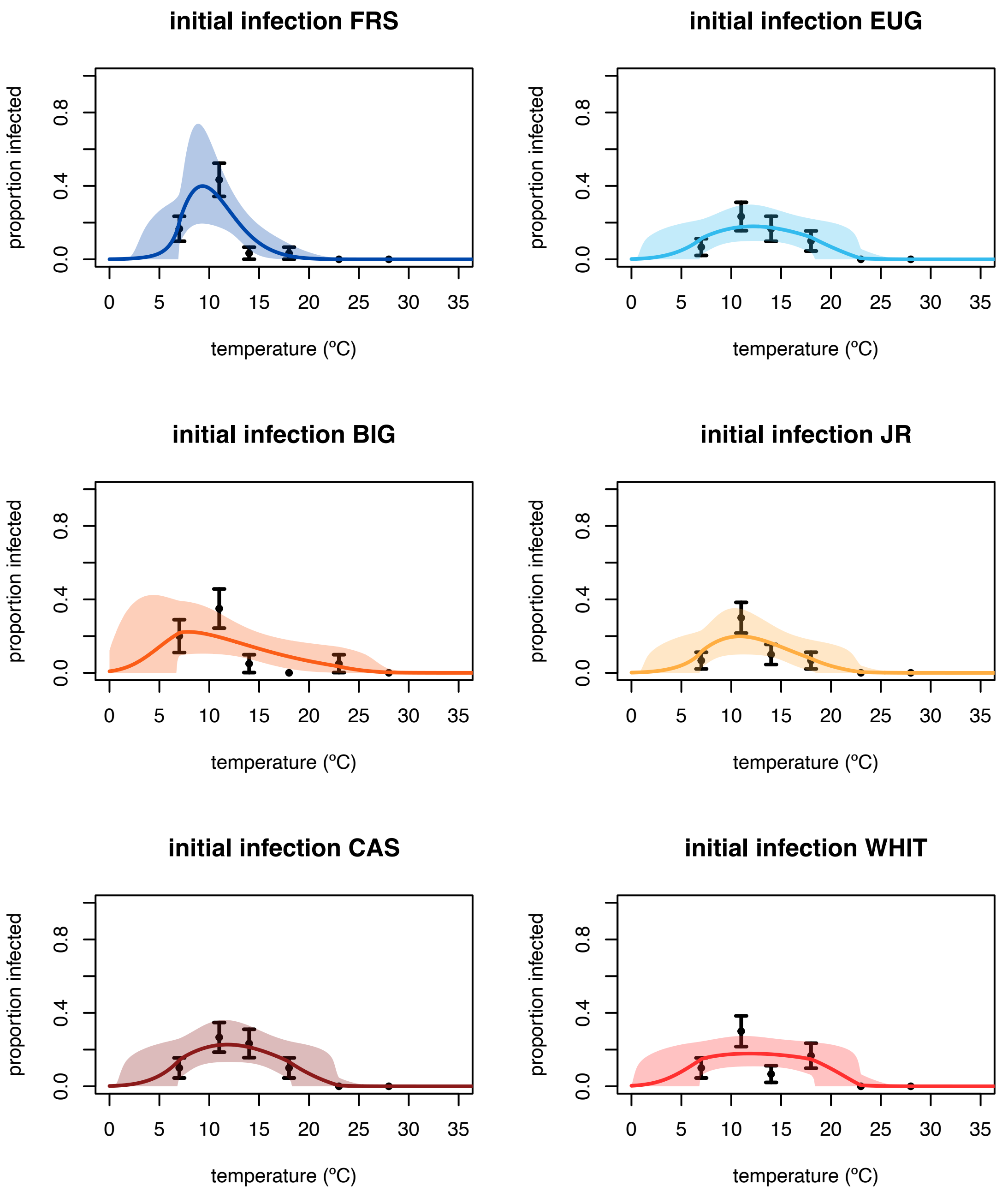


**Figure S2. Thermal responses of initial infection rates, representing primary infections that occurred prior to the first infected host death at each temperature.** Each panel shows the experimental data and TPC for one host population. Points show the mean proportion of larvae infected and error bars show +/- 1 SE, lines show the mean estimated thermal response, and shading shows the 95% CI for the TPC. Populations from cold climates are shown in the top row, intermediate climates in the middle row, and warm climates in the bottom row.

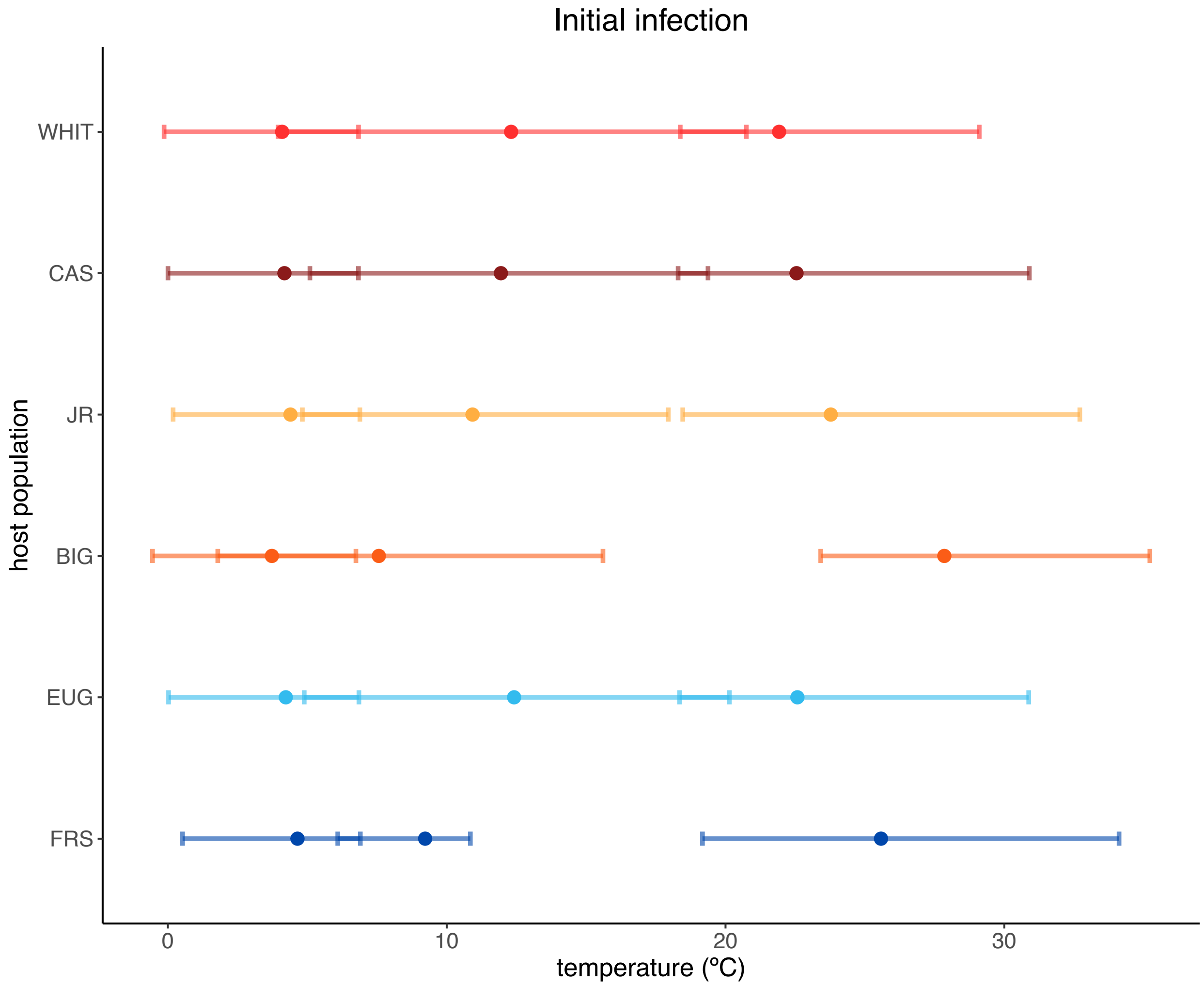


**Figure S3. Mean estimates and 95% credible intervals for initial infection $T_{min}$, $T_{opt}$, and $T_{max}$ for each experimental host population.** Host populations are ordered on the y-axis from the coldest source climate (bottom) to the warmest source climate (top). Colors correspond to host population IDs.

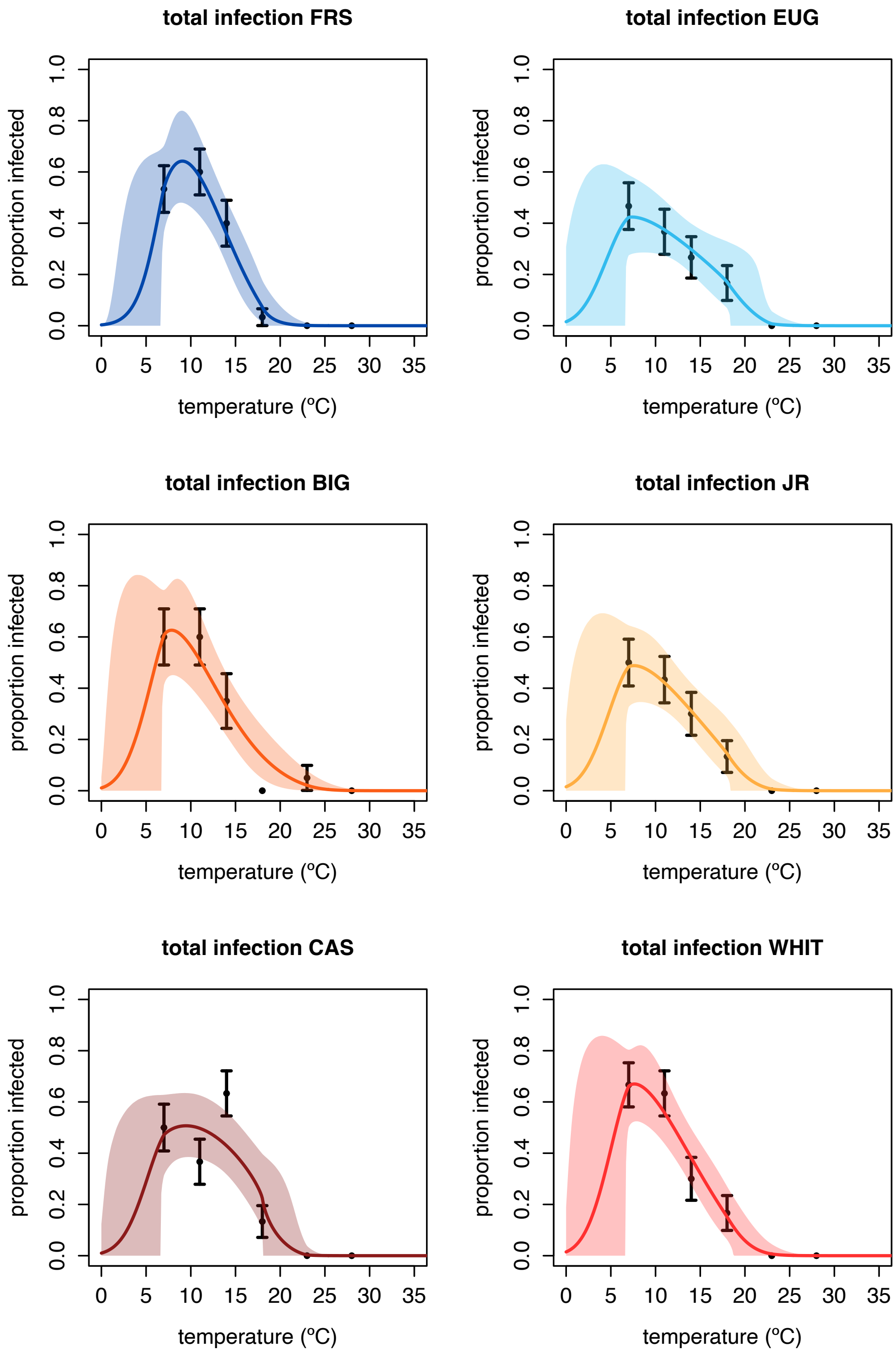


**Figure S4. Thermal responses of total infection rates for each population over the course of the entire experiment, representing both primary infections and those transmitted from killed hosts.** Each panel shows the experimental data and TPC for one host population. Points show the mean proportion of larvae infected and error bars show +/- 1 SE, lines show the mean estimated thermal response, and shading shows the 95% CI for the TPC. Populations from cold climates are shown in the top row, intermediate climates in the middle row, and warm climates in the bottom row.

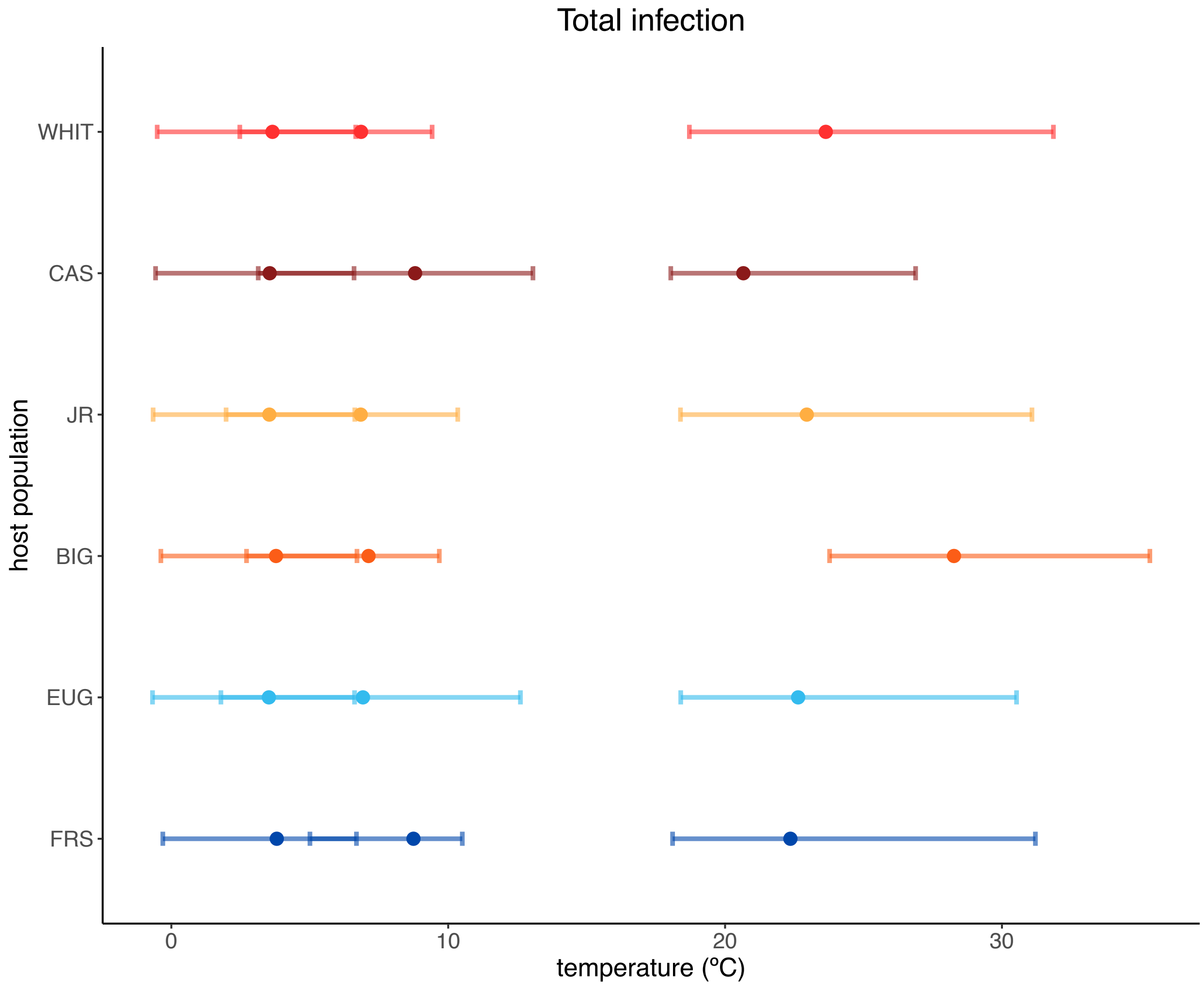


**Figure S5. Mean estimates and 95% credible intervals for total infection $T_{min}$, $T_{opt}$, and $T_{max}$ for each experimental host population.** Host populations are ordered on the y-axis from the coldest source climate (bottom) to the warmest source climate (top). Colors correspond to host population IDs.

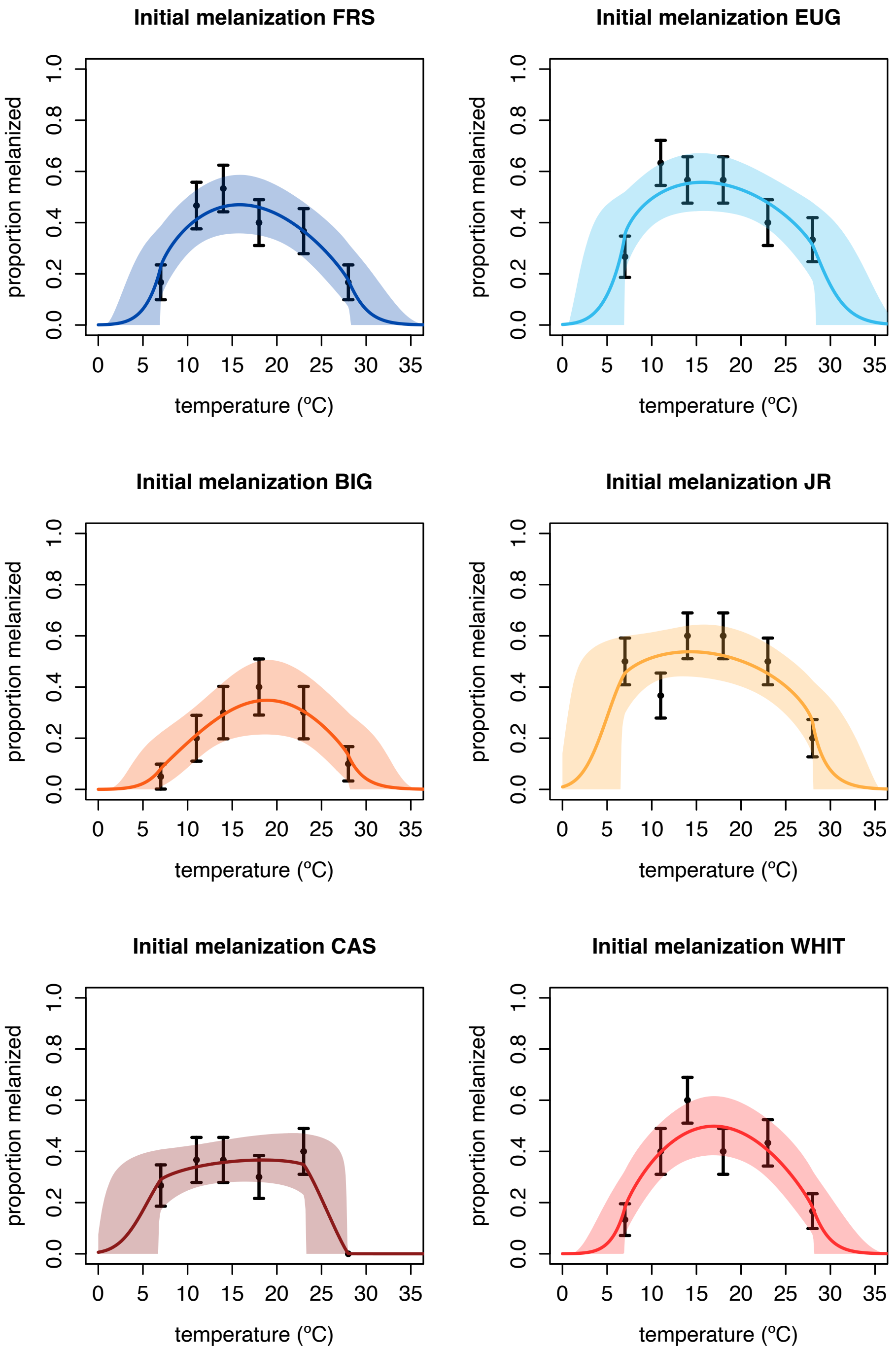


**Figure S6. Thermal responses of initial parasite melanization rates, representing responses to attack by parasites prior to the first infected host death at each temperature.** Each panel shows the experimental data and TPC for one host population. Points show the mean proportion of larvae infected and error bars show +/- 1 SE, lines show the mean estimated thermal response, and shading shows the 95% CI for the TPC. Populations from cold climates are shown in the top row, intermediate climates in the middle row, and warm climates in the bottom row.

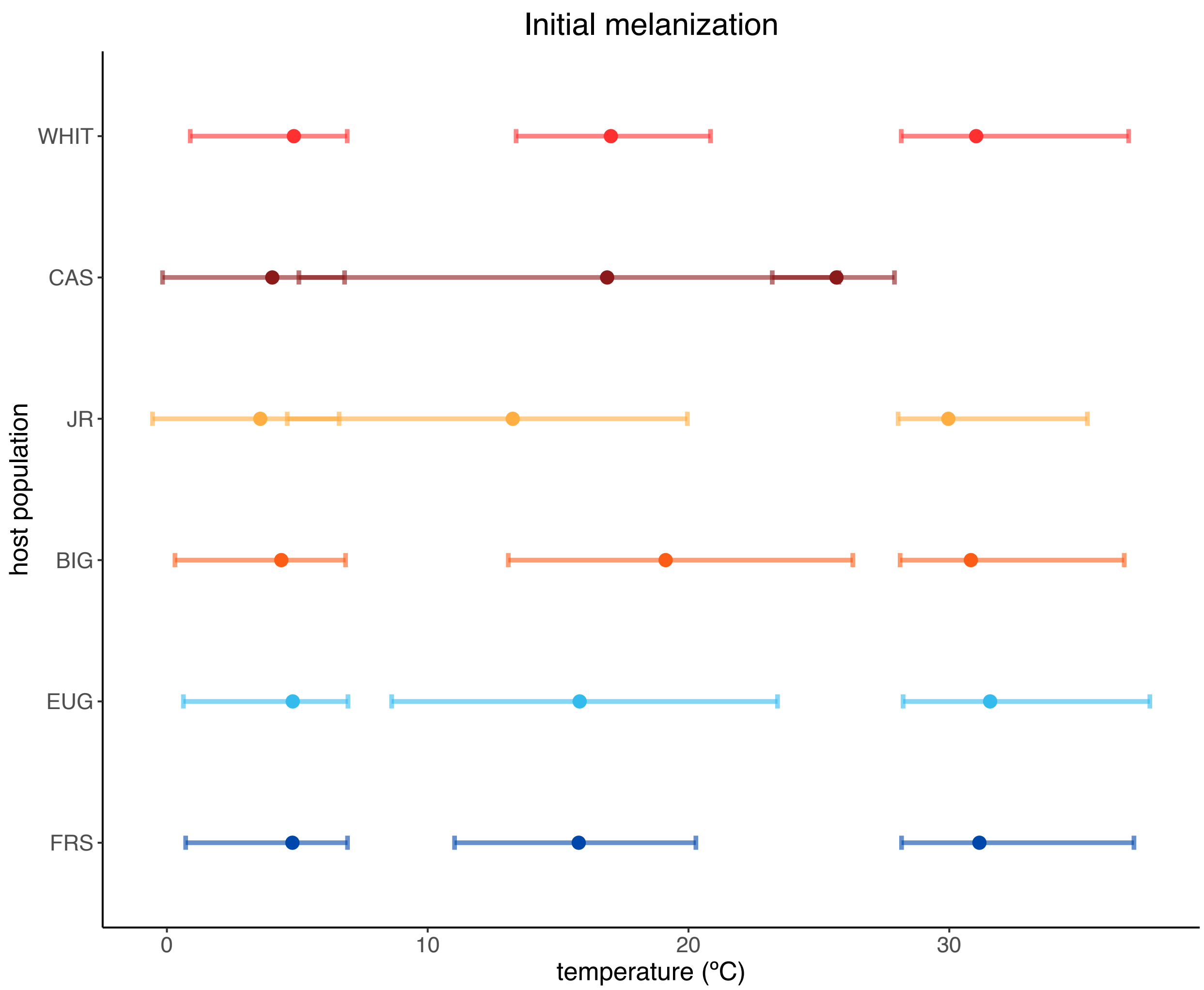


**Figure S7. Mean estimates and 95% credible intervals for initial melanization $T_{min}$, $T_{opt}$, and $T_{max}$ for each experimental host population.** Host populations are ordered on the y-axis from the coldest source climate (bottom) to the warmest source climate (top). Colors correspond to host population IDs.

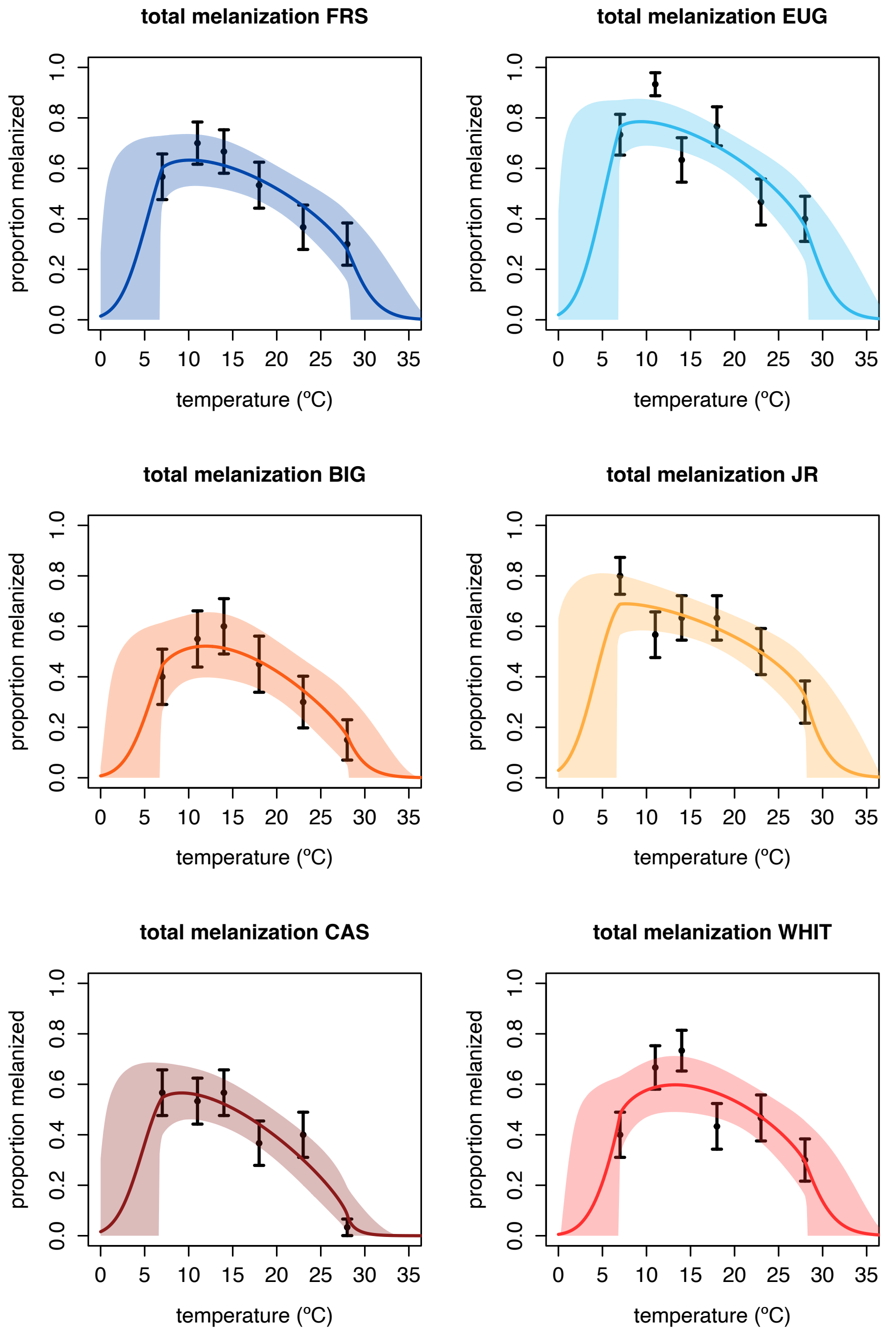


**Figure S8. Thermal responses of total parasite melanization rates for each population over the course of the entire experiment, representing responses to both initial parasite attack and later attack by parasites released by killed hosts.** Each panel shows the experimental data and TPC for one host population. Points show the mean proportion of larvae with melanization spots and error bars show +/- 1 SE, lines show the mean estimated thermal response, and shading shows the 95% CI for the TPC. Populations from cold climates are shown in the top row, intermediate climates in the middle row, and warm climates in the bottom row.

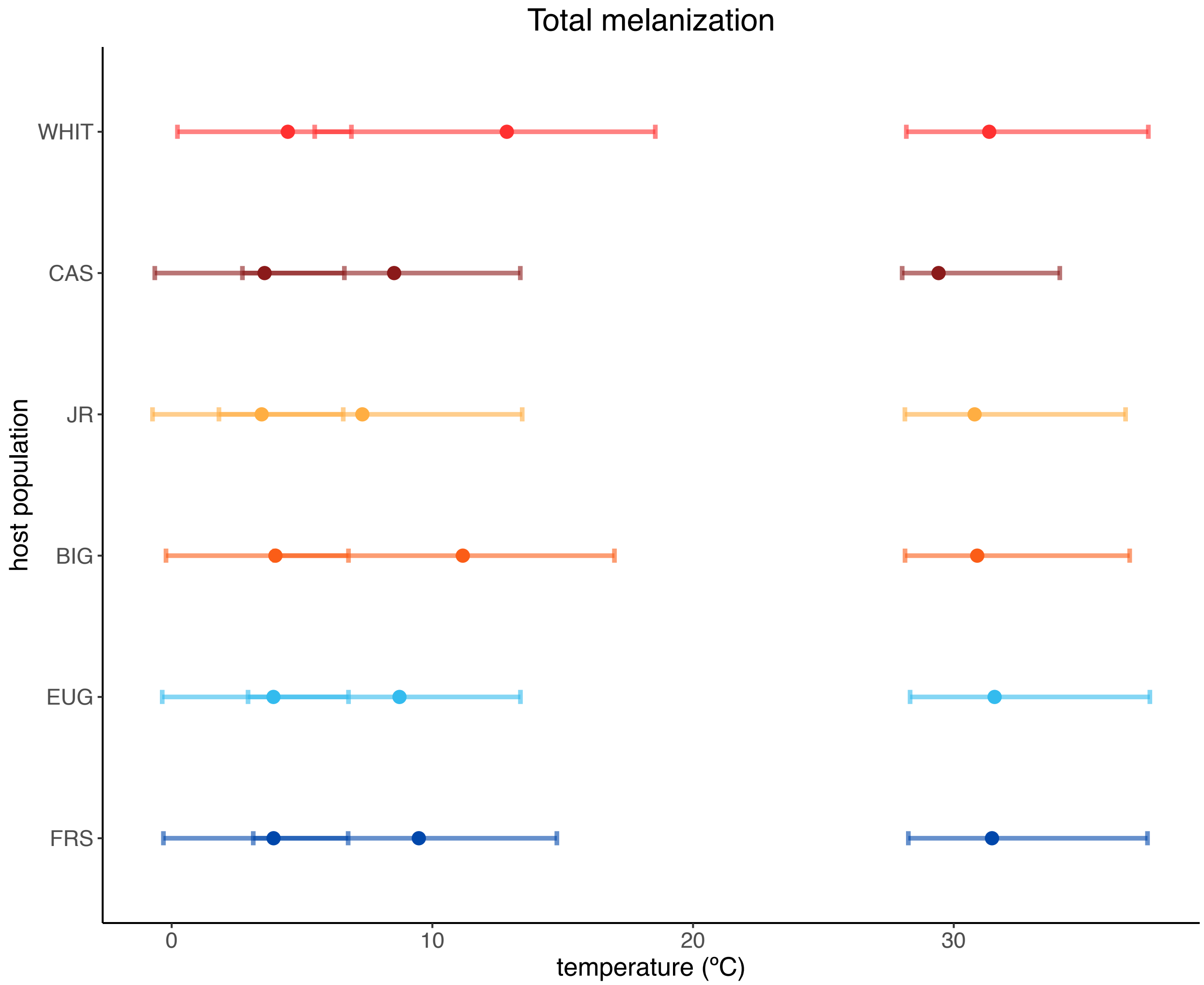


**Figure S9. Mean estimates and 95% credible intervals for total melanization $T_{min}$, $T_{opt}$, and $T_{max}$ for each experimental host population.** Host populations are ordered on the y-axis from the coldest source climate (bottom) to the warmest source climate (top). Colors correspond to host population IDs.

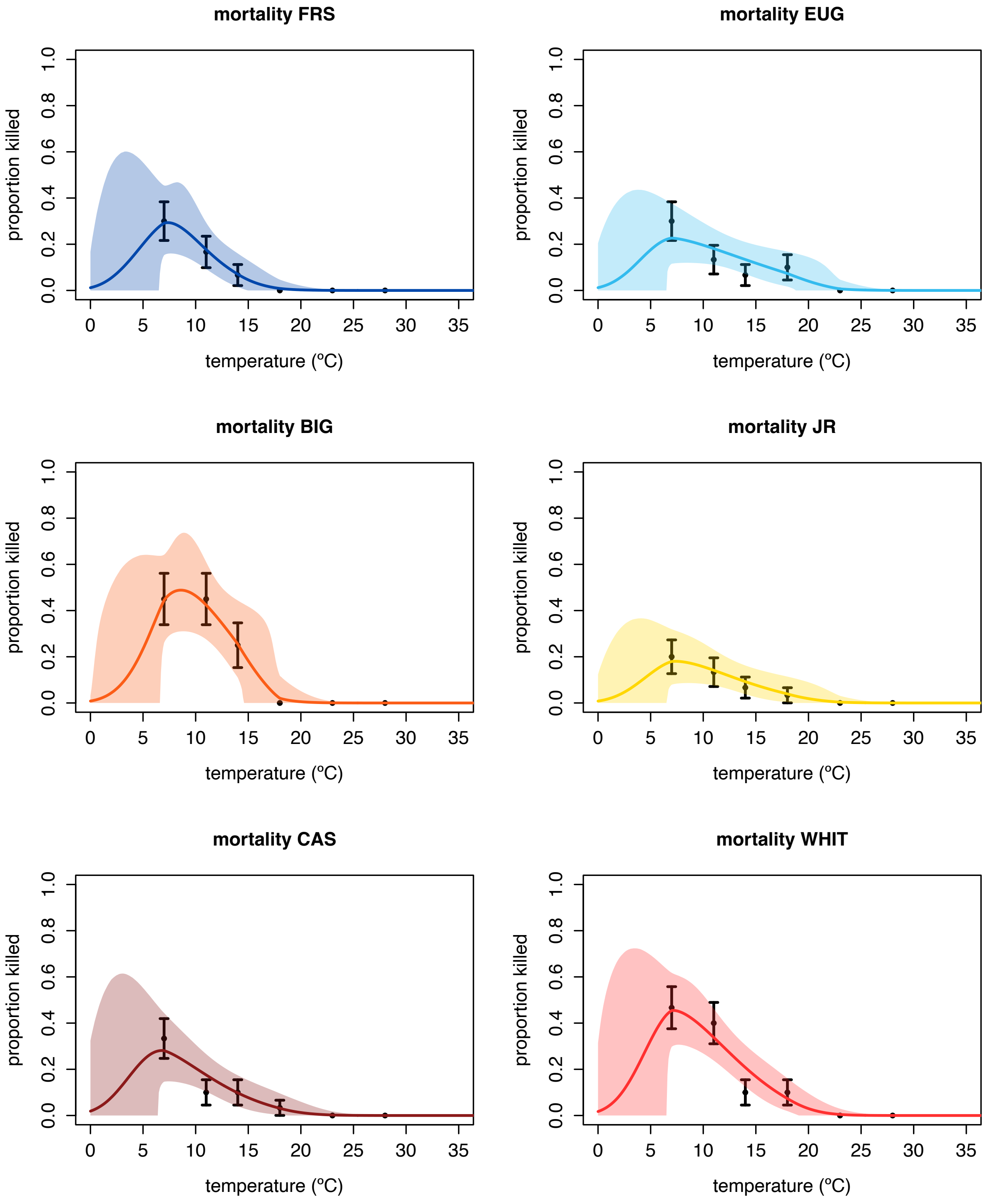


**Figure S10. Thermal responses of mortality of infected hosts, for each population over the course of the entire experiment.** Each panel shows the experimental data and TPC for one host population. Points show the mean proportion of larvae with melanization spots and error bars show +/- 1 SE, lines show the mean estimated thermal response, and shading shows the 95% CI for the TPC. Populations from cold climates are shown in the top row, intermediate climates in the middle row, and warm climates in the bottom row.

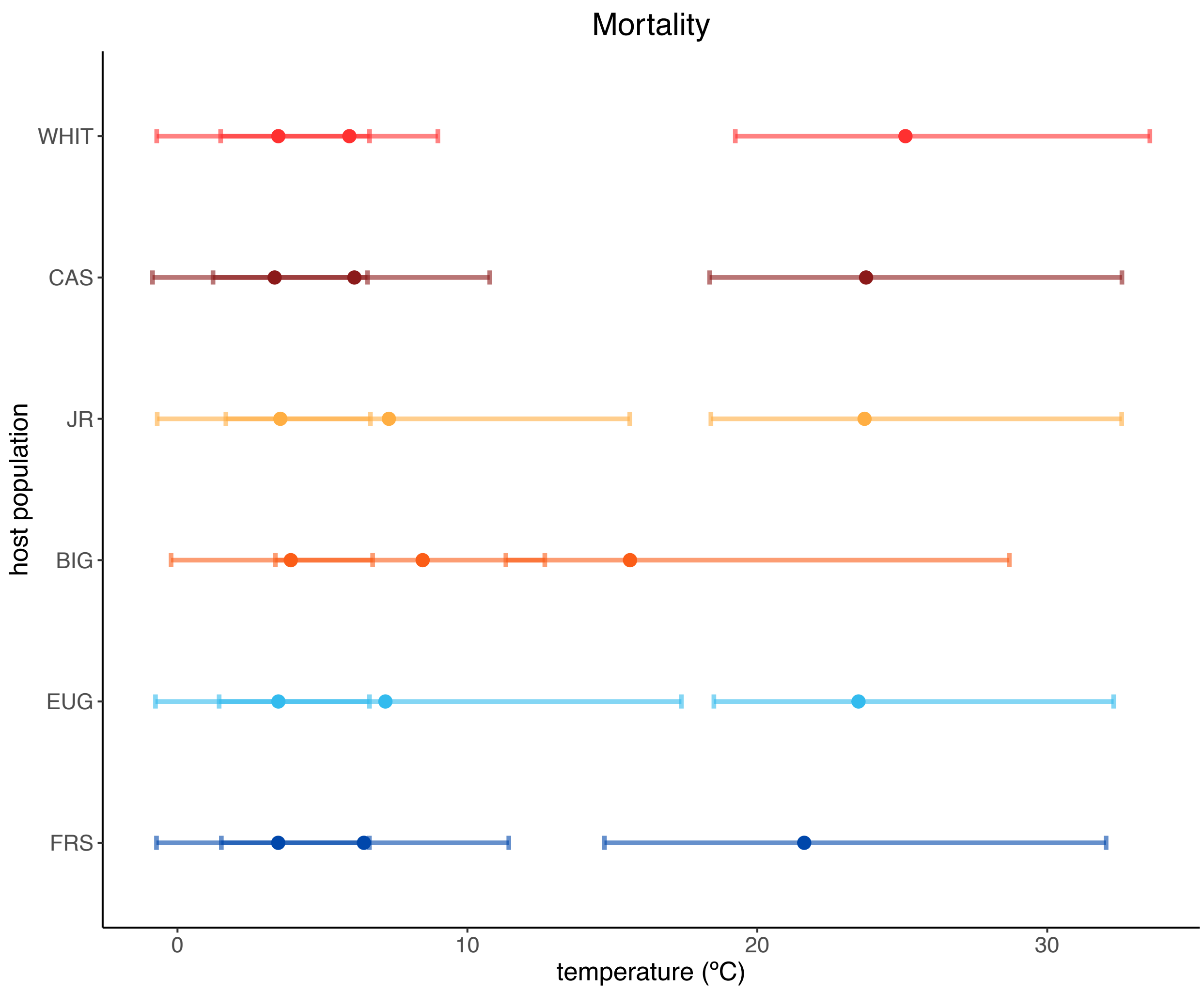


**Figure S11. Mean estimates and 95% credible intervals for mortality $T_{min}$, $T_{opt}$, and $T_{max}$ for each experimental host population.** Host populations are ordered on the y-axis from the coldest source climate (bottom) to the warmest source climate (top). Colors correspond to host population IDs.

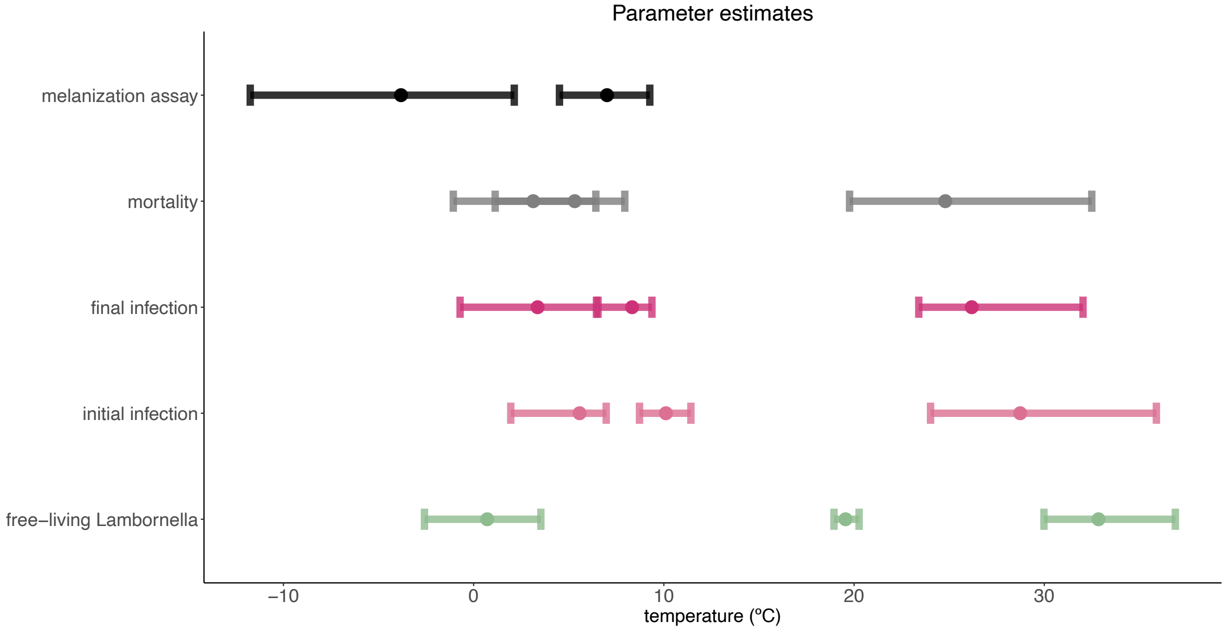


**Figure S12. Point estimates and credible intervals for TPC parameters varied among components and outcomes of the host – parasite interaction.** Mean estimates and 95% CIs for $T_{min}$ and $T_{half}$ are shown for the melanization assay TPC; estimates for $T_{min}$, $T_{opt}$, and $T_{max}$ are shown for mortality, infection, and free-living *Lambornella* growth rate TPCs.

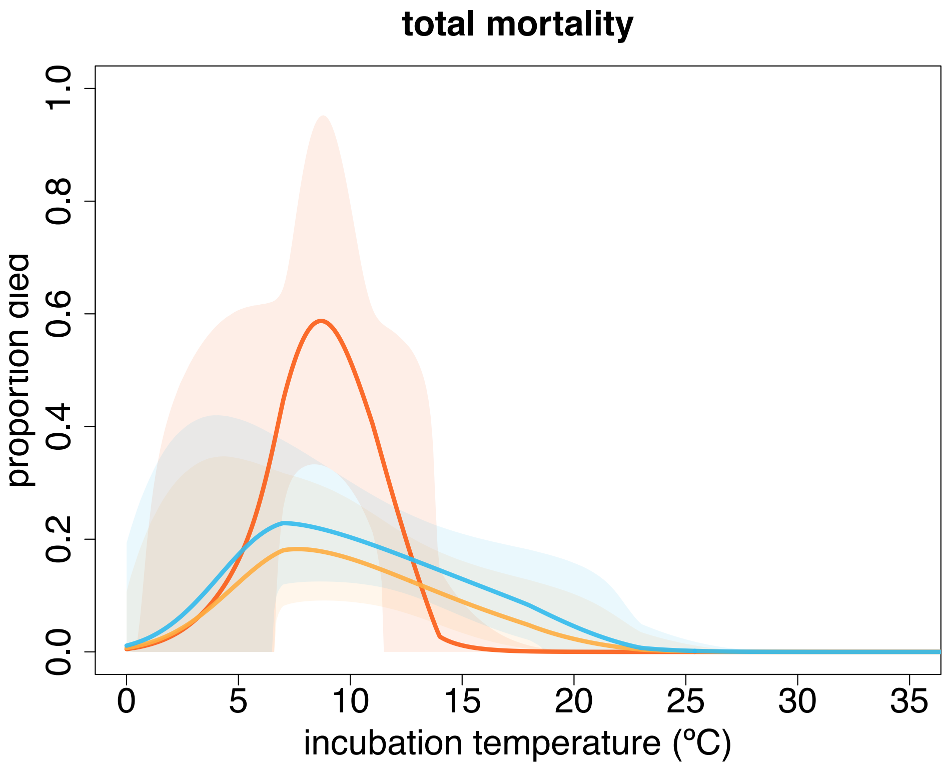


**Figure S13. Mortality rates near 10ºC were higher in the intermediate climate host population from Big Sur, CA (BIG; dark orange) compared to both the other intermediate climate population from Woodside, CA (JR; light orange) and to the cold climate population from Eugene, OR (EUG; light blue).** Lines show the mean estimated mortality TPCs and shading 95% CIs.

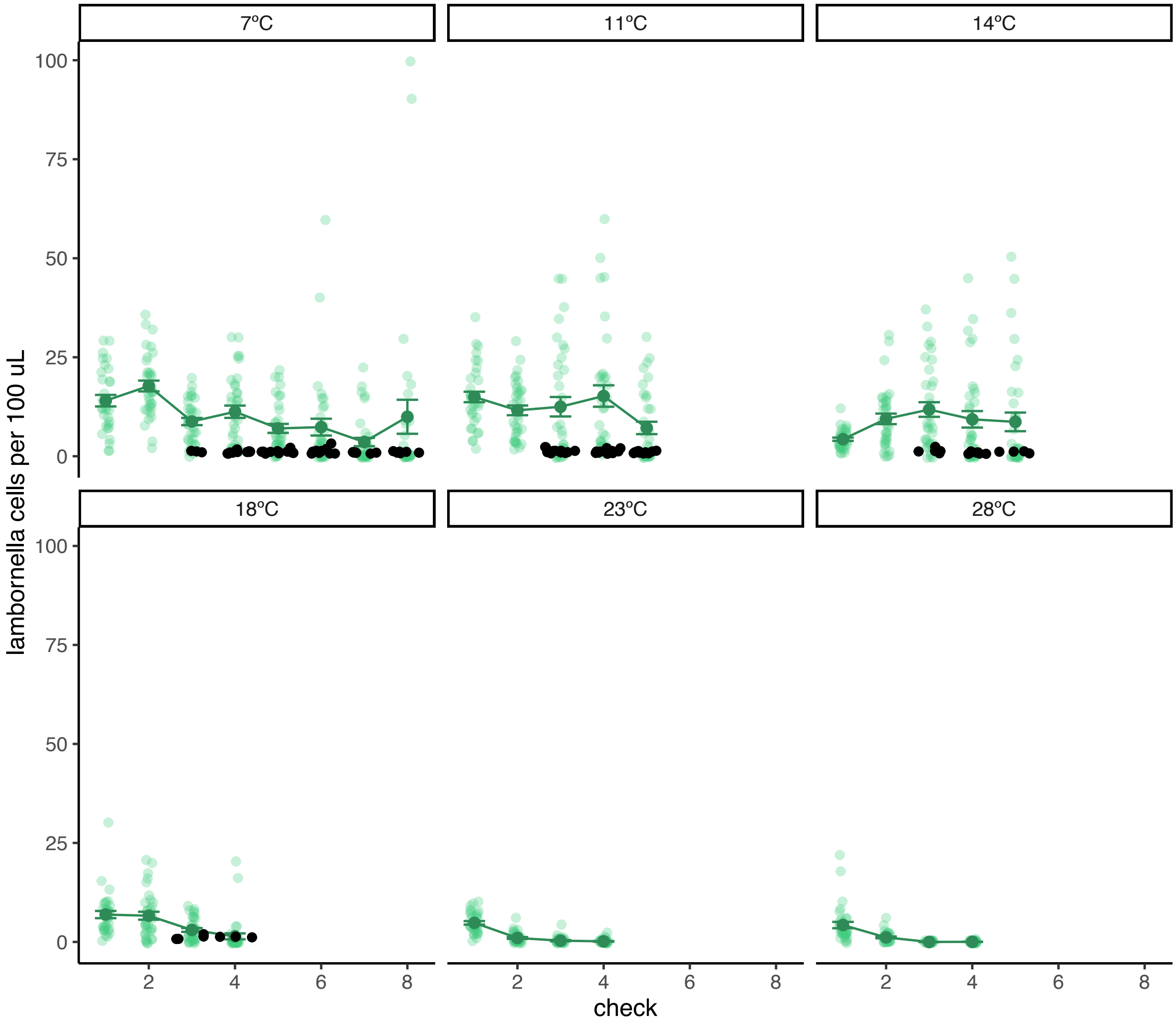


**Figure S14. At colder temperatures, *Lambornella clarki* cell density within microcosms was maintained as infected host deaths released new parasites under conditions suitable for *L. clarki* replication.** For each incubation temperature (panels), dark green points show the mean number of *L. clarki* cells per 100 uL liquid, light green points show counts for individual microcosms, and black points show observations of infected host deaths within microcosms.

**Tables**

**Table S1. Experimental mosquito and parasite population information. Rainy season mean temperature was calculated from PRISM data for 1991 – 2020.**

| Population ID | Role in experiment | Location | Coordinates | Rainy season mean temperature (ºC) |
|---|---|---|---|---|
| FRS | host | Shaver Lake, Fresno County, CA | 37.113, -119.311 | 5.3 |
| EUG | host | Eugene, Lane County, OR | 44.015, -123.09 | 6.5 |
| JR | host | Woodside, San Mateo County, CA | 37.407, -122.24 | 11.8 |
| BIG | host | Big Sur, Monterey County, CA | 36.014, -121.41 | 10.8 |
| CAS | host | San Juan Capistrano, Orange County, CA | 33.537, -117.55 | 15.1 |
| WHIT | host | Trabuco Canyon, Orange County, CA | 33.69, -117.65 | 14.8 |
| Marin | parasite | Novato, Marin County, CA | 38.130, -122.54 | 10.6 |
| PAR | melanization assay | Paso Robles, San Luis Obispo County, CA | 35.455, -120.69 | 10.9 |

**Table S2.** Frequency of monitoring for infection, parasite melanization, and mosquito survival at each incubation temperature in the infection experiment.

| Temperature | Microscope check interval | Number of microscope checks, ending with first pupation |
|---:|---:|:---:|
| 7ºC | 7 days | 8 |
| 11ºC | 7 days | 5 |
| 14ºC | 4 days | 5 |
| 18ºC | 4 days | 4 |
| 23ºC | 3 days | 4 |
| 28ºC | 2 days | 4 |

**Table S3**. Posterior distribution summaries for initial infection TPCs for each experimental host population. From left to right, columns list the mean parameter estimate, the lower and upper bounds of the 95% confidence intervals, the $\hat{R}$ convergence statistic, and the effective sample size, $N_{eff}$.

| Parameter | Population | Mean | 2.5% | 97.5% | $\hat{R}$ | $N_{eff}$ |
|---|---|---|---|---|---|---|
| $T_{opt}$ | FRS | 9.2 | 6.1 | 10.9 | 1.001 | 450000 |
| | EUG | 12.4 | 4.9 | 20.1 | 1.001 | 450000 |
| | BIG | 7.6 | 1.8 | 15.7 | 1.001 | 270000 |
| | JR | 10.9 | 4.8 | 17.9 | 1.001 | 450000 |
| | CAS | 11.9 | 5.1 | 19.4 | 1.001 | 450000 |
| | WHIT | 12.3 | 3.9 | 20.7 | 1.001 | 360000 |
| $T_{min}$ | FRS | 4.7 | 0.549 | 6.9 | 1.001 | 68000 |
| | EUG | 4.2 | 0.036 | 6.9 | 1.001 | 450000 |
| | BIG | 3.7 | -0.573 | 6.7 | 1.001 | 380000 |
| | JR | 4.4 | 0.186 | 6.9 | 1.001 | 160000 |
| | CAS | 4.2 | 0.001 | 6.8 | 1.001 | 240000 |
| | WHIT | 4.1 | -0.149 | 6.8 | 1.001 | 450000 |
| $T_{max}$ | FRS | 25.6 | 19.2 | 34.2 | 1.001 | 120000 |
| | EUG | 22.6 | 18.4 | 30.9 | 1.001 | 210000 |
| | BIG | 27.8 | 23.4 | 35.2 | 1.001 | 190000 |
| | JR | 23.8 | 18.5 | 32.7 | 1.001 | 450000 |
| | CAS | 22.6 | 18.3 | 30.9 | 1.001 | 450000 |
| | WHIT | 21.9 | 18.4 | 29.1 | 1.001 | 450000 |
| $\alpha$ | FRS | 0.220 | 0.064 | 0.426 | 1.001 | 84000 |
| | EUG | 0.458 | 0.062 | 0.938 | 1.001 | 360000 |
| | BIG | 0.162 | 0.006 | 0.553 | 1.001 | 360000 |
| | JR | 0.347 | 0.057 | 0.826 | 1.001 | 450000 |
| | CAS | 0.433 | 0.069 | 0.906 | 1.001 | 450000 |
| | WHIT | 0.470 | 0.037 | 0.962 | 1.001 | 420000 |
| $\beta$ | FRS | 0.780 | 0.574 | 0.936 | 1.001 | 79000 |
| | EUG | 0.542 | 0.062 | 0.938 | 1.001 | 290000 |
| | BIG | 0.838 | 0.447 | 0.994 | 1.001 | 120000 |
| | JR | 0.653 | 0.174 | 0.943 | 1.001 | 220000 |
| | CAS | 0.567 | 0.094 | 0.931 | 1.001 | 450000 |
| | WHIT | 0.530 | 0.038 | 0.963 | 1.001 | 350000 |
| $r_{max}$ | FRS | 0.414 | 0.207 | 0.756 | 1.001 | 120000 |
| | EUG | 0.191 | 0.105 | 0.315 | 1.001 | 340000 |
| | BIG | 0.249 | 0.110 | 0.483 | 1.001 | 110000 |
| | JR | 0.210 | 0.105 | 0.368 | 1.001 | 420000 |
| | CAS | 0.238 | 0.138 | 0.377 | 1.001 | 240000 |
| | WHIT | 0.189 | 0.113 | 0.295 | 1.001 | 320000 |

**Table S4.** Posterior distribution summaries for total infection TPCs for each experimental host population. From left to right, columns list the mean parameter estimate, the lower and upper bounds of the 95% confidence intervals, the $\hat{R}$ convergence statistic, and the effective sample size, $N_{eff}$.

| Parameter | Population | Mean | 2.5% | 97.5% | $\hat{R}$ | $N_{eff}$ |
|---|---|---|---|---|---|---|
| | FRS | 8.8 | 5.0 | 10.5 | 1.001 | 380000 |
| | EUG | 6.9 | 1.8 | 12.5 | 1.001 | 82000 |
| $T_{opt}$ | BIG | 7.1 | 2.7 | 9.7 | 1.001 | 150000 |
| | JR | 6.8 | 2.0 | 10.4 | 1.001 | 170000 |
| | CAS | 8.8 | 3.1 | 13.0 | 1.001 | 450000 |
| | WHIT | 6.9 | 2.5 | 9.4 | 1.001 | 450000 |
| | FRS | 3.8 | -0.307 | 6.7 | 1.001 | 450000 |
| | EUG | 3.5 | -0.679 | 6.6 | 1.001 | 450000 |
| $T_{min}$ | BIG | 3.8 | -0.384 | 6.7 | 1.001 | 280000 |
| | JR | 3.5 | -0.638 | 6.6 | 1.001 | 250000 |
| | CAS | 3.6 | -0.579 | 6.6 | 1.001 | 450000 |
| | WHIT | 3.7 | -0.506 | 6.7 | 1.001 | 290000 |
| | FRS | 22.3 | 18.1 | 31.2 | 1.001 | 39000 |
| | EUG | 22.6 | 18.4 | 30.5 | 1.001 | 130000 |
| $T_{max}$ | BIG | 28.3 | 23.8 | 35.4 | 1.001 | 450000 |
| | JR | 23.0 | 18.4 | 31.1 | 1.001 | 440000 |
| | CAS | 20.7 | 18.0 | 26.8 | 1.001 | 100000 |
| | WHIT | 23.6 | 18.7 | 31.9 | 1.001 | 84000 |
| | FRS | 0.267 | 0.067 | 0.486 | 1.001 | 450000 |
| | EUG | 0.179 | 0.007 | 0.543 | 1.001 | 280000 |
| $\alpha$ | BIG | 0.134 | 0.011 | 0.297 | 1.001 | 98000 |
| | JR | 0.170 | 0.009 | 0.425 | 1.001 | 440000 |
| | CAS | 0.307 | 0.024 | 0.646 | 1.001 | 310000 |
| | WHIT | 0.157 | 0.012 | 0.356 | 1.001 | 220000 |
| | FRS | 0.733 | 0.514 | 0.933 | 1.001 | 370000 |
| | EUG | 0.821 | 0.457 | 0.993 | 1.001 | 110000 |
| $\beta$ | BIG | 0.866 | 0.703 | 0.989 | 1.001 | 55000 |
| | JR | 0.830 | 0.575 | 0.991 | 1.001 | 450000 |
| | CAS | 0.693 | 0.354 | 0.976 | 1.001 | 450000 |
| | WHIT | 0.843 | 0.644 | 0.988 | 1.001 | 210000 |
| | FRS | 0.659 | 0.497 | 0.852 | 1.001 | 72000 |
| | EUG | 0.451 | 0.293 | 0.670 | 1.001 | 200000 |
| $r_{max}$ | BIG | 0.672 | 0.473 | 0.909 | 1.001 | 350000 |
| | JR | 0.520 | 0.357 | 0.737 | 1.001 | 230000 |
| | CAS | 0.528 | 0.395 | 0.671 | 1.001 | 450000 |
| | WHIT | 0.707 | 0.542 | 0.905 | 1.001 | 450000 |

**Table S5.** Posterior distribution summaries for initial melanization TPCs for each experimental host population. From left to right, columns list the mean parameter estimate, the lower and upper bounds of the 95% confidence intervals, the $\hat{R}$ convergence statistic, and the effective sample size, $N_{eff}$.

| Parameter | Population | Mean | 2.5% | 97.5% | $\hat{R}$ | $N_{eff}$ |
|---|---|---|---|---|---|---|
| $T_{opt}$ | FRS | 15.8 | 11.3 | 20.0 | 1.001 | 450000 |
| | EUG | 15.6 | 8.6 | 21.9 | 1.001 | 280000 |
| | BIG | 19.1 | 14.0 | 25.7 | 1.001 | 450000 |
| | JR | 13.0 | 4.0 | 19.5 | 1.001 | 190000 |
| | CAS | 18.2 | 3.7 | 26.7 | 1.001 | 220000 |
| | WHIT | 17.0 | 13.5 | 20.7 | 1.001 | 450000 |
| $T_{min}$ | FRS | 4.8 | 0.656 | 6.9 | 1.001 | 450000 |
| | EUG | 4.8 | 0.608 | 6.9 | 1.001 | 420000 |
| | BIG | 4.3 | 0.222 | 6.8 | 1.001 | 170000 |
| | JR | 3.5 | -0.625 | 6.6 | 1.001 | 230000 |
| | CAS | 3.9 | -0.278 | 6.8 | 1.001 | 450000 |
| | WHIT | 4.8 | 0.794 | 6.9 | 1.001 | 450000 |
| $T_{max}$ | FRS | 31.4 | 28.2 | 37.4 | 1.001 | 260000 |
| | EUG | 31.9 | 28.3 | 38.0 | 1.001 | 430000 |
| | BIG | 30.9 | 28.1 | 36.8 | 1.001 | 270000 |
| | JR | 30.2 | 28.1 | 35.6 | 1.001 | 160000 |
| | CAS | 25.7 | 23.2 | 27.9 | 1.001 | 420000 |
| | WHIT | 31.2 | 28.2 | 37.2 | 1.001 | 240000 |
| $\alpha$ | FRS | 0.415 | 0.200 | 0.641 | 1.001 | 450000 |
| | EUG | 0.402 | 0.133 | 0.699 | 1.001 | 250000 |
| | BIG | 0.559 | 0.313 | 0.836 | 1.001 | 450000 |
| | JR | 0.356 | 0.024 | 0.651 | 1.001 | 120000 |
| | CAS | 0.656 | 0.010 | 0.995 | 1.001 | 230000 |
| | WHIT | 0.464 | 0.270 | 0.673 | 1.001 | 450000 |
| $\beta$ | FRS | 0.413 | 0.236 | 0.738 | 1.001 | 450000 |
| | EUG | 0.525 | 0.284 | 1.0 | 1.001 | 450000 |
| | BIG | 0.349 | 0.186 | 0.711 | 1.001 | 60000 |
| | JR | 0.574 | 0.288 | 1.0 | 1.001 | 150000 |
| | CAS | 0.729 | 0.213 | 1.6 | 1.001 | 450000 |
| | WHIT | 0.383 | 0.227 | 0.645 | 1.001 | 310000 |
| $r_{max}$ | FRS | 0.478 | 0.366 | 0.595 | 1.001 | 450000 |
| | EUG | 0.566 | 0.455 | 0.679 | 1.001 | 450000 |
| | BIG | 0.359 | 0.224 | 0.517 | 1.001 | 110000 |
| | JR | 0.551 | 0.452 | 0.656 | 1.001 | 290000 |
| | CAS | 0.380 | 0.291 | 0.487 | 1.001 | 450000 |
| | WHIT | 0.506 | 0.392 | 0.624 | 1.001 | 450000 |

**Table S6.** Posterior distribution summaries for total melanization TPCs for each experimental host population. From left to right, columns list the mean parameter estimate, the lower and upper bounds of the 95% confidence intervals, the $\hat{R}$ convergence statistic, and the effective sample size, $N_{eff}$.

| Parameter | Population | Mean | 2.5% | 97.5% | $\hat{R}$ | $N_{eff}$ |
|---|---|---|---|---|---|---|
| $T_{opt}$ | FRS | 9.3 | 2.9 | 14.4 | 1.001 | 320000 |
| | EUG | 8.6 | 2.8 | 13.2 | 1.001 | 210000 |
| | BIG | 11.0 | 3.9 | 16.5 | 1.001 | 450000 |
| | JR | 7.0 | 1.7 | 12.8 | 1.001 | 120000 |
| | CAS | 8.5 | 2.8 | 13.3 | 1.001 | 310000 |
| | WHIT | 12.6 | 5.0 | 17.7 | 1.001 | 450000 |
| $T_{min}$ | FRS | 3.9 | -0.285 | 6.8 | 1.001 | 360000 |
| | EUG | 3.9 | -0.314 | 6.8 | 1.001 | 450000 |
| | BIG | 4.0 | -0.224 | 6.8 | 1.001 | 450000 |
| | JR | 3.5 | -0.697 | 6.6 | 1.001 | 450000 |
| | CAS | 3.6 | -0.625 | 6.6 | 1.001 | 270000 |
| | WHIT | 4.5 | 0.232 | 6.9 | 1.001 | 220000 |
| $T_{max}$ | FRS | 31.7 | 28.3 | 37.6 | 1.001 | 450000 |
| | EUG | 31.7 | 28.4 | 37.7 | 1.001 | 450000 |
| | BIG | 31.1 | 28.2 | 36.9 | 1.001 | 450000 |
| | JR | 30.9 | 28.2 | 36.7 | 1.001 | 380000 |
| | CAS | 29.5 | 28.0 | 34.2 | 1.001 | 450000 |
| | WHIT | 31.7 | 28.3 | 37.8 | 1.001 | 450000 |
| $\alpha$ | FRS | 0.190 | 0.014 | 0.420 | 1.001 | 79000 |
| | EUG | 0.164 | 0.013 | 0.372 | 1.001 | 95000 |
| | BIG | 0.260 | 0.025 | 0.510 | 1.001 | 450000 |
| | JR | 0.125 | 0.006 | 0.370 | 1.001 | 100000 |
| | CAS | 0.189 | 0.016 | 0.416 | 1.001 | 93000 |
| | WHIT | 0.300 | 0.040 | 0.550 | 1.001 | 310000 |
| $\beta$ | FRS | 0.454 | 0.196 | 0.816 | 1.001 | 210000 |
| | EUG | 0.459 | 0.196 | 0.784 | 1.001 | 86000 |
| | BIG | 0.424 | 0.211 | 0.793 | 1.001 | 400000 |
| | JR | 0.455 | 0.135 | 0.870 | 1.001 | 110000 |
| | CAS | 0.374 | 0.162 | 0.599 | 1.001 | 130000 |
| | WHIT | 0.517 | 0.275 | 0.965 | 1.001 | 450000 |
| $r_{max}$ | FRS | 0.647 | 0.540 | 0.757 | 1.001 | 400000 |
| | EUG | 0.799 | 0.700 | 0.894 | 1.001 | 420000 |
| | BIG | 0.537 | 0.408 | 0.675 | 1.001 | 450000 |
| | JR | 0.703 | 0.593 | 0.827 | 1.001 | 450000 |
| | CAS | 0.584 | 0.472 | 0.710 | 1.001 | 450000 |
| | WHIT | 0.608 | 0.500 | 0.720 | 1.001 | 450000 |

**Table S7.** Posterior distribution summaries for mortality TPCs for each experimental host population. From left to right, columns list the mean parameter estimate, the lower and upper bounds of the 95% confidence intervals, the $\hat{R}$ convergence statistic, and the effective sample size, $N_{eff}$.

| Parameter | Population | Mean | 2.5% | 97.5% | $\hat{R}$ | $N_{eff}$ |
|---|---|---|---|---|---|---|
| $T_{opt}$ | FRS | 6.5 | 1.7 | 10.0 | 1.001 | 95000 |
| | EUG | 6.7 | 1.5 | 15.1 | 1.001 | 300000 |
| | BIG | 8.5 | 3.4 | 12.8 | 1.001 | 450000 |
| | JR | 7.2 | 1.9 | 12.7 | 1.001 | 260000 |
| | CAS | 6.2 | 1.5 | 10.2 | 1.001 | 430000 |
| | WHIT | 6.1 | 1.7 | 9.1 | 1.001 | 450000 |
| $T_{min}$ | FRS | 3.4 | -0.800 | 6.6 | 1.001 | 250000 |
| | EUG | 3.4 | -0.843 | 6.6 | 1.001 | 450000 |
| | BIG | 3.7 | -0.341 | 6.7 | 1.001 | 110000 |
| | JR | 3.5 | -0.733 | 6.6 | 1.001 | 450000 |
| | CAS | 3.3 | -0.918 | 6.5 | 1.001 | 240000 |
| | WHIT | 3.4 | -0.795 | 6.6 | 1.001 | 400000 |
| $T_{max}$ | FRS | 21.8 | 14.9 | 32.0 | 1.001 | 250000 |
| | EUG | 23.8 | 18.6 | 32.4 | 1.001 | 200000 |
| | BIG | 17.4 | 11.5 | 30.3 | 1.001 | 12000 |
| | JR | 23.9 | 18.5 | 32.6 | 1.001 | 270000 |
| | CAS | 23.7 | 18.4 | 32.5 | 1.001 | 300000 |
| | WHIT | 25.1 | 19.3 | 33.4 | 1.001 | 200000 |
| $\alpha$ | FRS | 0.177 | 0.013 | 0.503 | 1.001 | 300000 |
| | EUG | 0.166 | 0.008 | 0.678 | 1.001 | 130000 |
| | BIG | 0.391 | 0.026 | 0.973 | 1.001 | 230000 |
| | JR | 0.185 | 0.012 | 0.553 | 1.001 | 96000 |
| | CAS | 0.144 | 0.009 | 0.406 | 1.001 | 190000 |
| | WHIT | 0.125 | 0.011 | 0.312 | 1.001 | 99000 |
| $\beta$ | FRS | 0.230 | 0.069 | 0.718 | 1.001 | 190000 |
| | EUG | 0.328 | 0.078 | 1.0 | 1.001 | 200000 |
| | BIG | 0.358 | 0.064 | 1.2 | 1.001 | 11000 |
| | JR | 0.276 | 0.081 | 0.815 | 1.001 | 74000 |
| | CAS | 0.237 | 0.070 | 0.594 | 1.001 | 450000 |
| | WHIT | 0.182 | 0.069 | 0.362 | 1.001 | 67000 |
| $r_{max}$ | FRS | 0.329 | 0.167 | 0.622 | 1.001 | 210000 |
| | EUG | 0.250 | 0.131 | 0.457 | 1.001 | 130000 |
| | BIG | 0.616 | 0.357 | 0.966 | 1.001 | 28000 |
| | JR | 0.204 | 0.097 | 0.387 | 1.001 | 450000 |
| | CAS | 0.309 | 0.163 | 0.563 | 1.001 | 450000 |
| | WHIT | 0.504 | 0.319 | 0.795 | 1.001 | 450000 |

**Table S8.** Posterior distribution summaries for TPCs fit to data combined across all experimental populations in the infection experiment. From left to right, columns list the mean parameter estimate, the lower and upper bounds of the 95% confidence intervals, the $\hat{R}$ convergence statistic, and the effective sample size, $N_{eff}$.

| Outcome | Parameter | Mean | 2.5% | 97.5% | $\hat{R}$ | $N_{eff}$ |
|---|---|---|---|---|---|---|
| initial infection | $T_{opt}$ | 10.0 | 8.7 | 11.3 | 1.001 | 95000 |
| | $T_{min}$ | 5.6 | 2.0 | 7.0 | 1.001 | 90000 |
| | $T_{max}$ | 29.0 | 24.2 | 36.1 | 1.001 | 450000 |
| | $\alpha$ | 0.188 | 0.076 | 0.345 | 1.001 | 76000 |
| | $\beta$ | 0.161 | 0.103 | 0.242 | 1.001 | 260000 |
| | $r_{max}$ | 0.277 | 0.207 | 0.373 | 1.001 | 81000 |
| total infection | $T_{opt}$ | 8.4 | 6.8 | 9.4 | 1.001 | 100000 |
| | $T_{min}$ | 3.3 | -0.723 | 6.4 | 1.001 | 89000 |
| | $T_{max}$ | 26.3 | 23.4 | 32.2 | 1.001 | 120000 |
| | $\alpha$ | 0.216 | 0.088 | 0.348 | 1.001 | 84000 |
| | $\beta$ | 0.200 | 0.136 | 0.268 | 1.001 | 320000 |
| | $r_{max}$ | 0.597 | 0.530 | 0.675 | 1.001 | 140000 |
| initial melanization | $T_{opt}$ | 16.2 | 14.1 | 18.4 | 1.001 | 250000 |
| | $T_{min}$ | 4.4 | 0.552 | 6.7 | 1.001 | 230000 |
| | $T_{max}$ | 30.5 | 28.2 | 35.4 | 1.001 | 84000 |
| | $\alpha$ | 0.454 | 0.308 | 0.598 | 1.001 | 130000 |
| | $\beta$ | 0.420 | 0.272 | 0.634 | 1.001 | 150000 |
| | $r_{max}$ | 0.495 | 0.447 | 0.545 | 1.001 | 450000 |
| total melanization | $T_{opt}$ | 9.6 | 5.1 | 12.2 | 1.001 | 200000 |
| | $T_{min}$ | 4.2 | 0.040 | 6.8 | 1.001 | 250000 |
| | $T_{max}$ | 31.9 | 28.6 | 37.7 | 1.001 | 46000 |
| | $\alpha$ | 0.191 | 0.046 | 0.342 | 1.001 | 57000 |
| | $\beta$ | 0.423 | 0.269 | 0.612 | 1.001 | 29000 |
| | $r_{max}$ | 0.658 | 0.610 | 0.710 | 1.001 | 140000 |
| mortality | $T_{opt}$ | 3.9 | -0.098 | 6.9 | 1.001 | 560000 |
| | $T_{min}$ | 2.9 | -1.3 | 6.3 | 1.001 | 46000 |
| | $T_{max}$ | 36.8 | 31.9 | 42.9 | 1.001 | 140000 |
| | $\alpha$ | 0.029 | 0.002 | 0.093 | 1.002 | 3700 |
| | $\beta$ | 0.095 | 0.027 | 0.178 | 1.002 | 3700 |
| | $r_{max}$ | 0.378 | 0.275 | 0.546 | 1.001 | 35000 |

**Table S9.** Posterior distribution summaries for the free-living *Lambornella clarki* population growth rate from the parasite-only experiment. From left to right, columns list the mean parameter estimate, the lower and upper bounds of the 95% confidence intervals, the $\hat{R}$ convergence statistic, and the effective sample size, $N_{eff}$.

| Parameter | Mean | 2.5% | 97.5% | $\hat{R}$ | $N_{eff}$ |
|---|---|---|---|---|---|
| $T_{opt}$ | 19.5 | 18.9 | 20.2 | 1.001 | 22000 |
| $T_{min}$ | 0.736 | -2.6 | 3.5 | 1.001 | 26000 |
| $T_{max}$ | 32.9 | 30.0 | 367.0 | 1.001 | 17000 |
| $\alpha$ | 0.586 | 0.513 | 0.654 | 1.001 | 18000 |
| $\beta$ | 0.290 | 0.229 | 0.368 | 1.001 | 23000 |
| $r_{max}$ | 1.7 | 1.6 | 1.7 | 1.001 | 140000 |
| $\sigma$ | 0.091 | 0.070 | 0.119 | 1.001 | 450000 |

**Table S10.** Posterior distribution summaries for the mechanically stimulated mosquito melanization immune response TPC. From left to right, columns list the mean parameter estimate, the lower and upper bounds of the 95% confidence intervals, the $\hat{R}$ convergence statistic, and the effective sample size, $N_{eff}$.

| Parameter | Mean | 2.5% | 97.5% | $\hat{R}$ | $N_{eff}$ |
|---|---|---|---|---|---|
| $T_{half}$ | 7.0 | 4.5 | 9.3 | 1.001 | 230000 |
| $T_{min}$ | -3.8 | -11.8 | 2.1 | 1.001 | 150000 |
| $\alpha$ | 9.7 | 2.4 | 27.5 | 1.001 | 14000 |
| $s$ | 0.909 | 0.024 | 2.4 | 1.001 | 10000 |
